# Novel phylogenomic inference and ‘Out of Asia’ biogeography of cobras, coral snakes, and their allies

**DOI:** 10.1101/2024.05.17.594737

**Authors:** Jeffrey L. Weinell, Frank T. Burbrink, Sunandan Das, Rafe M. Brown

## Abstract

Estimation of evolutionary relationships among lineages that rapidly diversified can be challenging, and, in such instances, inaccurate or unresolved phylogenetic estimates can lead to erroneous conclusions regarding historical geographical ranges of lineages. One example underscoring this issue has been the historical challenge posed by untangling the biogeographic origin of elapoid snakes, which includes numerous dangerously venomous species as well as species not known to be dangerous to humans. The worldwide distribution of this lineage makes it an ideal group for testing hypotheses related to historical faunal exchanges among the many continents and other landmasses occupied by contemporary elapoid species. We developed a novel suite of genomic resources, included worldwide sampling, and inferred a robust estimate of evolutionary relationships, which we leveraged to quantitatively estimate geographical range evolution through the deep-time history of this remarkable radiation. Our phylogenetic and biogeographical estimates of historical ranges definitively reject a lingering former ‘Out of Africa’ hypothesis and support an ‘Out of Asia’ scenario involving multiple faunal exchanges between Asia, Africa, Australasia, the Americas, and Europe.

## 1. Introduction

How, why, and when geographic distributions of lineages have changed through time are primary questions motivating the conceptual unification of phylogenetic systematics, macroecology, and Earth history in the multidisciplinary field of modern biogeography. Characteristics of geography, climate, species interactions, and historical contingency have all been implicated in determining present day geographic ranges of species [1]. Tests of biogeographic hypotheses related to historical faunal exchanges require well-documented information about the geographic ranges of species, knowledge of Earth history and historical environmental changes, and robust analytical methods for inferring evolutionary relationships and historical geographic ranges of lineages. As a result of centuries of field-based exploration and biogeographical research [2–6], geographic ranges have been broadly characterized, at least at a continental resolution, for most known terrestrial vertebrate species [7,8], and evolutionary relationships have been inferred for many vertebrate lineages as a result of molecular phylogenetic studies over the last half century [9–15]. However, for groups within which little is known about species diversity, evolutionary relationships, or geographic ranges of lineages—commonly known as Linnean, Darwinian, and Wallacean shortfalls [16–19]—biogeographic origins are often unknown and can be hard to accurately estimate.

One such group is the snake superfamily Elapoidea, which includes over 700 species in nine families and is geographically distributed in tropical and subtropical regions on all the planet’s major continents and in marine habitats of Indian and Pacific oceans (figure 1) [20–22]. The group’s biogeographic origin, however, has remained uncertain. Earlier studies have supported an African origin for this lineage—implying its ancestral expansion ‘Out of Africa’—and highlighted its rapid diversification in Africa [21–29]. An African origin has also been supported for numerous additional vertebrate groups presently or historically distributed on multiple continents [30–37]. One significant barrier that has precluded robust quantitative biogeographical investigation of Elapoidea persists: a strongly supported, time-calibrated estimate of evolutionary relationships, including all extant major (family-rank level) lineages and sublineages, has remained unavailable.

**Figure 1.**
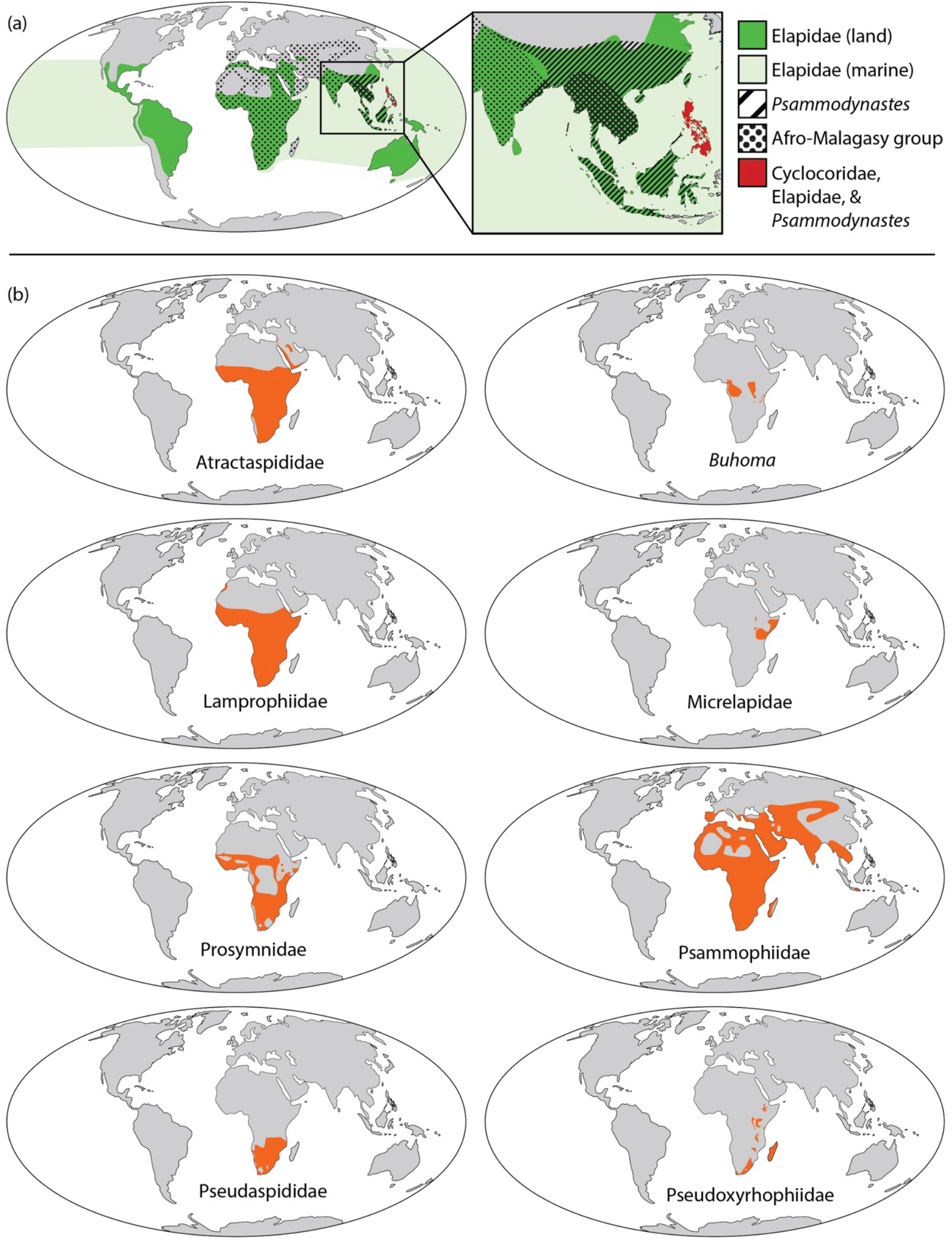
Geographic extents of major lineages of Elapoidea; (a) geographic distributions for Cyclocoridae (red), Elapidae (green or red), *Psammodynastes* (diagonal black stripes or red), and the group comprising non-elapid Afro-Malagasy lineages (black circle pattern); (b) geographic extent (orange) for each major lineage (family-rank or similar) of the Afro-Malagasy group.

Here, we used a targeted sequence capture approach to produce a phylogeny that samples all major elapoid lineages and outgroups. We used this phylogenomic tree and information from the fossil record of snakes to estimate a strongly supported, time-calibrated phylogenetic framework for implementation of modern biogeographical inference. We generated the most comprehensive biogeographical inference performed to date with respect to taxonomic coverage, number of loci, and the use of powerful, statistically coherent methods. Using our results of biogeographic analyses, we assess previous hypotheses related to historical faunal exchange and the ‘Out of Africa’ scenario supported by phylogenetic, biogeographic, and paleontological results of earlier studies [21,25,38]. Results supporting ancestral range expansions into or out of Africa before formation of a land bridge connection between this continent and Eurasia would necessarily indicate that such expansions likely resulted from historical oceanic dispersal events.

## 2. Materials and Methods

### 2.1. Sequence Capture Dataset

We sampled 66 individuals from 65 species in 52 genera and 22 families of snakes, from DNA isolated from ethanol-preserved tissue samples or from previous assembled genomes (electronic supplementary material, table S1). Tissues were collected by numerous individuals (electronic supplementary material) during field expeditions (1990–2018) and deposited in the tissue collections at the University of Kansas Biodiversity Institute, University of Texas at El Paso, or Villanova University (Villanova, PA).

We targeted 3128 loci and 1,517,011 nucleotides (nt) for capture with a MyBaits-20 custom RNA probe kit (Arbor BioSciences) containing 20,020 120nt probes and 2X probe tiling. Target loci included 1652 rapidly evolving exons (REEs), 907 UCEs, 328 ddRAD-like (*in silico* identification), 27 major histo-compatability (MHC), 119 vision-associated, and 95 scalation-associated loci (electronic supplementary material; Open Science Framework project krhx3). Targeted regions of REEs included 121–7501nt, one or more entire exons, and partial upstream, downstream, or both up- and downstream exon-flanking regions. Sequence capture libraries were dual-indexed and paired-end sequenced (150 bp inserts) on an Illumina HiSeq X sequencer (NovoGene).

We demultiplexed fastq files with “demuxbyname” from BBMap [39], trimmed adapters with FASTP [40], filtered contaminant reads by comparison to a database of common contaminants with BBMap [41,42], removed duplicate reads with “dedupe” tool in BBTools, and merged read pairs with BBMerge [43]. To *de novo* assemble reads into contigs, we used SPAdes 3.12 [44] with multiple k-mer values and dipSPAdes in SPAdes [45] for polymorphic read assembly. We masked low complexity and repeat regions with repeatMasker 4.1.0 [46]. To annotate contigs, we used BLASTn from BLAST+ 2.9 to match contigs against probe target reference sequences, filtered matches with bitscore <50, and binned contigs into disjoint clusters with R package ‘igraph’ and SeqKit 2 [47–49].

To obtain orthologous sequences from previously assembled genomes (electronic supplementary material, table S1) for each locus in our novel sequence capture dataset, we aligned novel sequences using MAFFT 7.310 [50] and generated a consensus sequence of the alignment result using R package ‘Biostrings’ 2.60.2 [51]; searched for matches to the consensus sequence in 20kb windows of genome assembly scaffolds using BLASTn [52]; pairwise aligned sequences with a strong match (bitscore >50); trimmed contiguous sites at ends of each pairwise alignment containing missing data for either sequence; retained the genome-derived sequence from each trimmed pairwise alignment; and removed gaps. We excluded loci, when multiple homologous regions were identified in the same genome assembly, and we considered sequences of included loci to be orthologous. To generate every single locus alignment used for phylogenetic estimation of gene trees, we aligned sequence capture and genome-derived orthologs using MAFFT 7.310, and trimmed each alignment down to the largest contiguous region containing 90% complete data. To generate our concatenation dataset, we concatenated single locus alignments using SeqKit 2 and R package ‘Biostrings’ 2.60.2 [49,51,53].

### 2.2. Sanger dataset

To estimate a chronogram with dense taxonomic sampling for our biogeographic analyses, we sampled genetic data for 450 individuals from 434 species of snakes in 379 genera and 24 families and non-nominate subfamilies. We compiled DNA sequences from GenBank (n = 428 individuals) and generated novel genetic data from ethanol-preserved tissue samples of 22 individuals of 21 species and genera (electronic supplementary material, table S2). Sampled loci included the mitochondrial protein-coding genes cytochrome b (CYTB), and five nuclear-encoded genes: Brain Derived Neurotrophic Factor (BDNF), oocyte maturation factor mos (C-MOS), 3’-Nucleotidase (NT3), Recombination Activating 1 (RAG1), and Recombination Activating 2 (RAG2). To extract and purify genomic DNA, we digested tissues using Proteinase K enzyme and used a Maxwell® Rapid Sample Concentrator Instrument with Maxwell® 16 Tissue DNA Purification Kit (Promega Corporation). We used 34 cycles polymerase chain reaction (PCR) with annealing temperature 49ºC (± 1–5ºC) and previously designed PCR and sequencing primers (electronic supplementary material, table S3) to amplify 1350 base pairs (bp) of CYTB and 580bp of C-MOS. Amplified products were visualized using gel electrophoresis on 1.5% agarose gels. Purification of PCR products, cycle sequencing, cycle sequencing clean-ups, and nucleotide sequence determination were conducted with standard GeneWiz protocols®. We *de novo* assembled and edited Sanger sequences using Geneious® 6.1 and Geneious Prime® 2021.2.2, aligned sequences separately for each locus with MAFFT 7.310 [50,54], and used Geneious Prime® and SeqKit 2 [49] to trim alignment ends, identify reading frames, filter out individuals with early stop codons in protein coding regions, and concatenate alignments.

### 2.3. Phylogenetic inference

To estimate ML gene trees from our single locus alignments and ML phylogeny from our concatenated sites alignment, we ran IQ-TREE 2.2 [55] with node support assessed from 1000 ultrafast bootstraps (UFBoot) [56]. For estimation of gene trees from single locus alignments, sites were partitioned into the disjoint set of protein-coding and non-coding regions determined by mapping protein-coding annotations from genome annotation files to our single locus alignments (electronic supplementary material; Open Science Framework, project krhx3), and we used option “-m TESTMERGE” of ModelFinder in IQ-TREE 2.1.3 to estimate the optimal partitioning scheme and to identify the best-fit substitution model for each partition. For phylogenetic estimation using our concatenated informative sites alignment, sites were unpartitioned and we used option “-m TEST” of ModelFinder in IQ-TREE 2.1.3 to identify the best substitution model. We considered UFBoot ≥95 and SH-aLRT ≥80 to be strong support for monophyly [55].

To infer the species tree with local posterior probability (PP) values of node support, we used ASTRAL-III 5.6.1 [57] and quartet trees sampled from our estimated gene trees with low support nodes (UFBoot ≤10) collapsed into polytomies; we considered PP >0.95 to be strong support for monophyly [57]. We used R packages ‘ape’ 5.6.1 [58] and ‘Quartet’ 1.2.2 [59] to estimate metrics of similarity and dissimilarity among gene, species, and concatenation tree topologies, including Robinson-Foulds Distance (RFD), Strict Joint Assertions (SJA), and Quartet Divergence (QDiv).

We inferred phylogenetic relationships using our Sanger sequence alignment and IQ-TREE 2.1.3 and optimal site-partitioning scheme estimated using ModelFinder implemented in IQ-TREE with option “-m TESTMERGEONLYNEW”. During phylogenetic inference, we used a highly resolved topological constraint tree congruent at the family-rank level with our inferred species tree and with additional topological constraints corresponding to a curated list of clades strongly supported in previous phylogenetic studies (electronic supplementary material; Open Science Framework project krhx3). To initiate tree optimization, we used a starting tree with topology generated by randomly resolving multichotomies in our constraint tree using function “multi2di” from R package ‘ape’ 5.6.1 [58]. For unconstrained nodes, we assessed node support from 1000 ultrafast bootstraps (UFBoot) and 1000 replicates for Shimodaira-Hasegawa (SH)-like approximate likelihood ratio testing [56,60].

### 2.4. Divergence time estimation

We estimated divergence times for our inferred “sequence capture species tree” and “Sanger ML tree” using RelTime-ML [61–63] in MEGA 11 [63]; an alignment with 10,000 sites randomly sampled without replacement from our sequence capture dataset (for analysis with inferred species tree) or the full Sanger sequence alignment (for analysis with inferred Sanger tree); and five node-calibration age constraints with lognormally distributed probability densities generated using information from the fossil record (electronic supplementary material, table S4). Calibrated nodes and confidence intervals (CIs) of calibrated node age probability densities included: Viperinae stem node (CI: 23.97–51.24 Ma); Elapidae stem node (CI: 26.77–54.04 Ma); Dipsadidae stem node (CI: 14.37–41.64 Ma); Colubroidea stem node (CI: 37.07–64.34 Ma); and Oxyuraninae stem node (CI: 11.87–39.14 Ma).

### 2.5. Historical hybridization and incomplete lineage sorting

To test for historical hybridizations, we estimated phylogenetic networks from gene concordance factors (gCFs) using SNAQ [64] implemented in the JULIA package ‘PhyloNetworks’ 0.14.3 [65,66]. We compared quartet trees sampled from our gene trees against the set of quartets obtained by quartet decomposition of the species tree to estimate gCFs, ran SNAQ ten times under each *a priori* maximum number of reticulations (which we varied from zero to ten), and retained the maximum pseudolikelihood network for each number of reticulations [64,67]. To estimate the optimal number of reticulations from the set of maximum pseudolikelihood networks, we used Djump and DDSE algorithms in R package ‘capushe’ 1.1.1 [68], which calculate slope heuristics considering model complexity and change in pseudolikelihood with each increase in number of reticulations [69]. We considered the optimal phylogenetic network to be the maximum pseudolikelihood network that had the outgroup lineage correctly inferred as the outgroup, and number of the reticulations equal to the inferred optimal number of reticulations. We considered historical hybridization to be supported if the optimal phylogenetic network contained a hybrid edge.

To assess if our phylogenetic results deviated from expectations of ILS, we estimated gene discordance factors (gDF1 and gDF2) and site discordance factors (sDF1 and sDF1) for internal nodes of our species tree and the phylogeny estimated from our concatenated-locus alignment, using IQ-TREE 2.1.3, our single-locus alignments, and quartets sampled from our estimated gene trees. We conducted χ^2^-tests of the hypothesis of no difference between sDF1 and sDF2, and no difference between gDF1 and gDF2, with significance considered at the 0.05 level [70,71]. Following the recommendation of Lanfear (2018), we binned putatively unlinked sites from our concatenated sites alignment by locus, bootstrap sampled one site per locus (100 bootstraps), and estimated sCF and sDFs from each bootstrap ‘unlinked sites alignment’ relative to our phylogenetic trees, and repeated χ^2^-tests on each bootstrapped sample to determine robustness. Significant χ^2^- test results were considered evidence against ILS-derived discordance [70], and, when combined with a low sCF or low gCF, as evidence suggesting ancestral hybridization.

### 2.6. Biogeographic inference

We inferred historical geographic ranges using BIOGEOBEARS 1.1.2 [72,73], our Sanger chronogram pruned to one tip per genus of superfamilies Colubroidea and Elapoidea, and a geographic distribution dataset [74] compiled from relevant literature sources and online databases [7,20,75–82]. We pruned all but one species lineage per genus and treated the geographic distribution of each retained lineage as equal to the total range of the corresponding genus.

We coded geographic distributions as either present or absent in six regions (electronic supplementary material, figure S1), including five physiogeographic land regions (Africa, Americas, Asia, Australasia, and Europe) and one marine region encompassing extant sea snake distributions (i.e., Indian and Pacific oceans). Within the “Africa” region, we included continental Africa, Madagascar, and most of the Arabian Plate in the Middle East. We considered continental North and South America and islands of the Caribbean, western Atlantic, and eastern Pacific oceans to comprise a single “Americas” region. The division between “Asia” and “Europe” was considered to occur approximately where the Turgai Strait likely separated these continents until 45–29 Ma [83,84]. We included islands of the Sunda Shelf and the Philippines within our concept of “Asia”. Within “Australasia”, we included continental Australia and Sahul Shelf and southwestern Pacific islands. We used these regions for biogeographic analyses because they are thought to have existed as distinct biogeographical and physiogeographical areas for most of the time hypothesized to span elapoid evolutionary history (⪅30 Ma).

During biogeographical analyses, we considered six geographic range evolution models: the (1) Dispersal-Extinction-Cladogenesis (DEC), (2) DEC+J, (3) DIVALIKE, (4) DIVALIKE+J, (5) BAYAREALIKE, and (6) BAYAREALIKE+J models, which differ in the types of range evolution processes that can occur during cladogenesis [72,73]. The BAYAREALIKE model does not allow range evolution to occur during cladogenesis; the DEC model allows vicariance or partial sympatric speciation if either daughter lineage occupies a single region. The DIVALIKE model allows vicariant speciation even if both daughter lineages have ranges spanning multiple regions, and sympatric speciation if the ancestor occupies a single area. The models DEC+J, DIVALIKE+J and BAYAREALIKE+J allow founder speciation, whereby one daughter lineage acquires a narrow range not occupied by the ancestor, and range evolution processes allowed by DEC, DIVALIKE and BAYAREALIKE models, respectively [72,73]. Ree & Sanmartín [85] warned against using statistical model selection to compare DEC and DEC+J models against other biogeographic models, whereas Matzke [86] considered such statistical model selection as valid. We considered the optimal biogeographic model to have the lowest Akaike Information Criterion (AIC) [87] during statistical model selection. To avoid over interpretive reliance on a single biogeographic estimate and implied evolutionary process, we compared ancestral geographic ranges estimated under each suboptimal biogeographic model to our results recovered under the optimal model.

## 3. Results

### 3.1. Sequence data

Using targeted sequence capture, we obtained new genetic data for 34 individuals from 32 species (electronic supplementary material, table S1), and our genetic datasets used for phylogenetic analyses included 3066 single-locus alignments and a concatenated-loci alignment with 66 individuals, 15,301,560 sites, and 77% total missing or ambiguous data (Open Science Framework, project krhx3). Number of loci sampled per individual ranged from 2041 (*Boa constrictor*) to 2717 (*Hologerrhum philippinum*). Single-locus alignments included 4– 66 sequences (median 59), 1–35 novel sequences (median 32), 528–58,675 sites (median 3697), 7–9733 informative sites (median 783), and 0.05–92.5% total missing or ambiguous data (median 65.12%); alignment length and percent missing data were both positively correlated with number of aligned sequences (*p* << 0.05; R^2^ =0.021 and 0.062, respectively).

Our Sanger genetic dataset included CYTB sampled for all individuals (n = 450; 1069 sites), and nuclear loci were sampled for subsets of individuals: BDNF (n = 24; 673 sites); C-MOS (n = 225; 588 sites); NT3 (n = 29; 516 sites); RAG1 (n = 33; 1000 sites); and RAG2 (n = 36; 714 sites). The concatenated sequence alignment contained 4560 sites and 69.3% missing or ambiguous data. Novel Sanger data included DNA sequences for 22 individuals of 21 species for loci CYTB (n = 22) and C-MOS (n = 21) and can be accessed in GenBank (electronic supplementary material, table S2).

### 3.2. Phylogenetic inference

Results from our ML phylogenetic analyses of individual loci in our sequence capture dataset included gene trees that were strongly supported at 0–100% of nodes (median 45.3%), and node support was significantly correlated with alignment length, number of aligned individuals, and number of informative sites (*p* <<0.05); 402 gene trees included all 66 individuals and had low to moderate topological variation (mean pairwise QDiv = 0.816; range = 0.408–0.987). Gene trees recovered Elapoidea as either monophyletic (51.8%), paraphyletic (n = 47.9%), or monophyly was indeterminable (n = 0.3%) due to limited sampling (i.e., one or no ingroup taxa sampled, or no outgroup sampled); Lamprophiidae: 11.5% monophyletic, 85.5% paraphyletic, 3% indeterminable; Elapidae: 81.7% monophyletic, 16.1% paraphyletic, 2.2% indeterminable; Cyclocoridae: 63.4% monophyletic, 33.1% paraphyletic, and 3.5% indeterminable; *Psammodynastes*: 70.1% monophyletic, 4.2% paraphyletic, 25.6% indeterminable; non-elapid Afro-Malagasy elapoids: 15.3% monophyletic, 82% paraphyletic, 2.7% indeterminable.

Our estimated species tree (figure 2*a*) and ML concatenation tree (figure 2*b*) were each strongly supported at most nodes (PP ≥0.95 at 94% of nodes of species tree; UFBoot ≥95 at 92% of nodes of concatenation tree) and had highly similar, but not identical, topologies (RFD = 20; QDiv =0.99). Our phylogenetic results strongly supported a clade composed of all non-elapid Afro-Malagasy elapoid lineages (figure 2), hereafter simply called the “Afro-Malagasy group”, composed of Atractaspididae, Lamprophiidae, Micrelapidae, Psammophiidae, Pseudaspididae, Pseudoxyrhophiidae, and Prosymnidae, and *Buhoma*.

**Figure 2.**
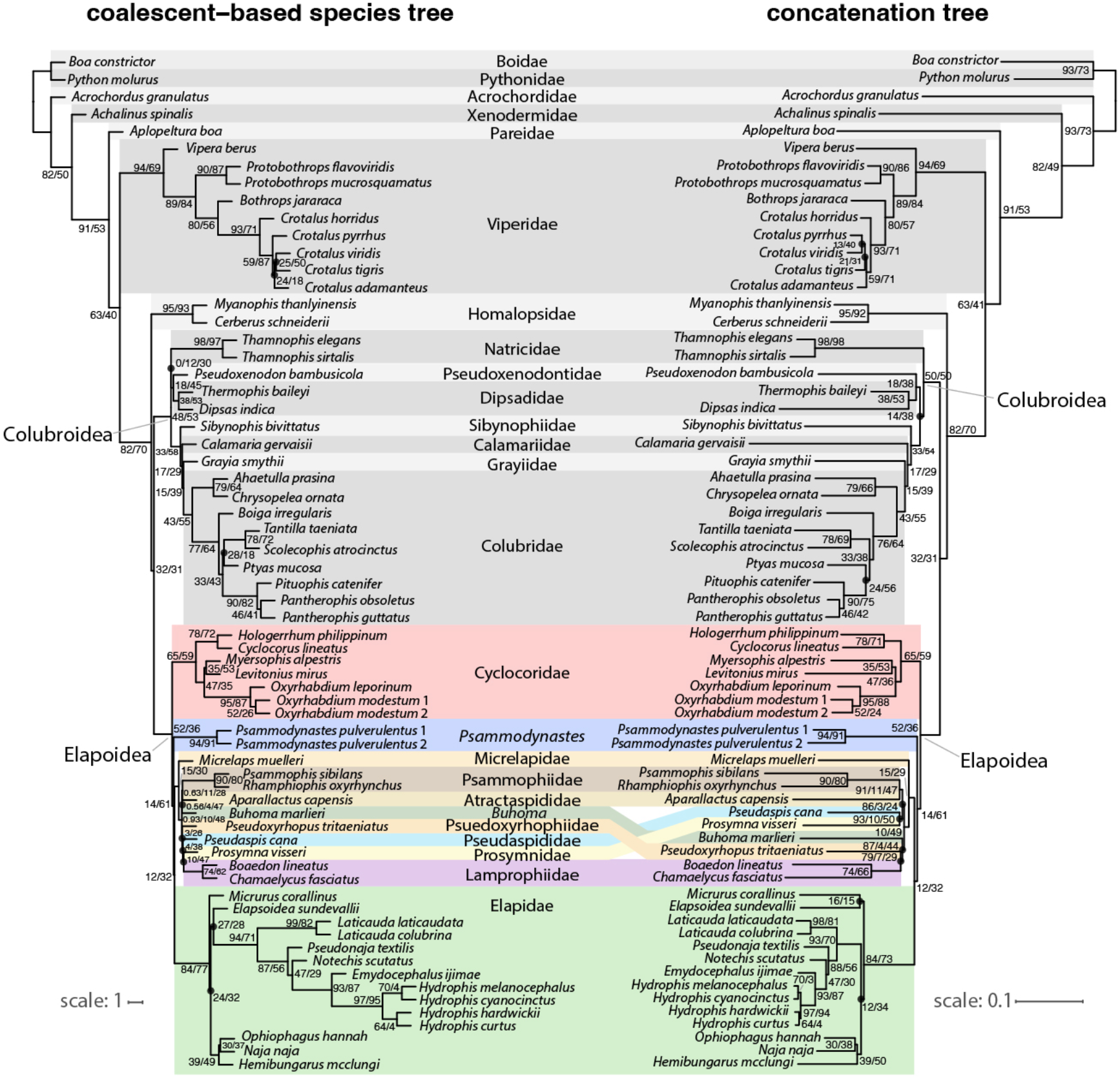
Coalescent-based species tree (left) and maximum likelihood concatenation tree (right). Black circles indicate groups unique to either the species tree or concatenation tree. Two node values indicate gene and site concordance factors, ‘gCF/sCF’, and strong support for group monophyly (LPP ≥ 0.95 for species tree; UFBoot ≥ 95 for concatenation tree); three node values indicate ‘LPP/gCF/sCF’ (species tree) or ‘UFBoot/gCF/sCF’ (concatenation tree). Branch lengths are in coalescent units (species tree) or substitutions per site (concatenation tree).

The species tree analysis inferred a sister relationship between Natricidae and the lineage comprising Dipsadidae and Pseudoxenodontidae, whereas the concatenation tree inferred the Dipsadidae-Pseudoxenodontidae and Natricidae lineages to be sequential outgroups of the clade comprising all other colubroid lineages. For the species tree and concatenation tree, inferred evolutionary relationships also differed among *Pantherophis*-*Pituophis, Ptyas*, and *Scolecophis-Tantilla* lineages (in Colubridae); among *Elapsoidea, Hemibungarus*-*Naja*-*Ophiophagus, Micrurus*, and Oxyuraninae lineages (in Elapidae); among *Crotalus adamanteus, C. pyrrhus, C. tigris*, and *C. viridis* (in Viperidae); among *Buhoma*, Atractaspididae, Lamprophiidae, Prosymnidae, Psammophiidae, Pseudaspididae, and Pseudoxyrhophiidae lineages (in Elapoidea). The optimal phylogenetic network inferred from our SNAQ analyses in ‘capushe’ was the network selected under the Djump algorithm, and did not include any reticulations (electronic supplementary material, table S5).

Gene trees were usually topologically consistent with the species tree and concatenation tree (mean QDiv = 0.82 between gene trees and the species tree; mean QDiv = 0.82 between gene trees and the concatenation tree). Low conflict among gene trees and each multilocus tree was also reflected in gene concordance and discordance factors inferred at nodes of the species and concatenation trees, such that gCF was usually higher than either of gDF1 and gDF2. For the species tree, gCF > gDF1 and gCF > gDF2 except at five nodes: (1) the most recent common ancestor (MRCA) of Lamprophiidae and Pseudoxyrhophiidae; (2) MRCA of Lamprophiidae and Pseudaspididae; (3) MRCA of *Crotalus tigris* and *C. viridis*; (4) MRCA of Dipsadidae and Natricidae; and (5) MRCA of Colubridae and Grayiidae. For the concatenation tree, gCF > gDF1 and gCF > gDF2 except at five nodes: (1) MRCA of *Crotalus pyrrhus* and *C. viridis*; (2) MRCA of *Hydrophis* and *Ophiophagus*; (3) MRCA of *Elapsoidea* and *Micrurus*; (4) MRCA of Colubridae and Grayiidae; and (5) MRCA of *Pantherophis, Pituophis*, and *Ptyas*. For both the species tree and concatenation tree, sDF1 and sDF2 significantly differed in 0.05% of 6400 χ^2^-tests (64 nodes; 100 bootstraps) and never significantly differed at the same node in more than one bootstrap, consistent with ILS-only discordance expectations; gDF1 and gDF2 significantly differed at 48% or 37% of nodes of the species tree or concatenation tree, respectively (electronic supplementary material, figure S2).

Our Sanger phylogeny, estimated from analysis of our Sanger sequence dataset, included 448 tips (individuals) sampled from 432 species and 378 genera, and was strongly supported (UFBoot ≥ 95) at 60% of nodes (n = 327) that were topologically unconstrained during phylogenetic analysis.

### 3.3. Divergence times

For lineages sampled in both our inferred species tree and Sanger tree, estimates of divergence times (means and limits of CIs) were generally older in the species tree than in the Sanger, especially at deeper nodes, even though node-calibrations were identical and tree topologies were consistent. Age estimates were similar between these trees at crown nodes of Colubroidea (species tree CI: 36.39–48.81 Ma; Sanger tree CI: 31.13– 44.33 Ma), Elapoidea (species tree CI: 35.63–45.92 Ma; Sanger tree CI: 28.94–41.24 Ma), and for colubroid and elapoid sublineages (figures 3–4). The greater number of lineages sampled in the Sanger tree compared to the species tree, fewer loci sampled and inclusion of mitochondrial loci in the Sanger dataset (versus thousands of loci, all from the nuclear genome, sampled in the sequence capture dataset used to estimate divergence times for the species tree) likely explain the differences in divergence times estimated for the species tree and Sanger tree. After pruning our time-calibrated Sanger phylogeny to include one tip per genus of superfamilies Colubroidea and Elapoidea, the chronogram used for biogeographic inference contained 311 tips.

**Figure 3.**
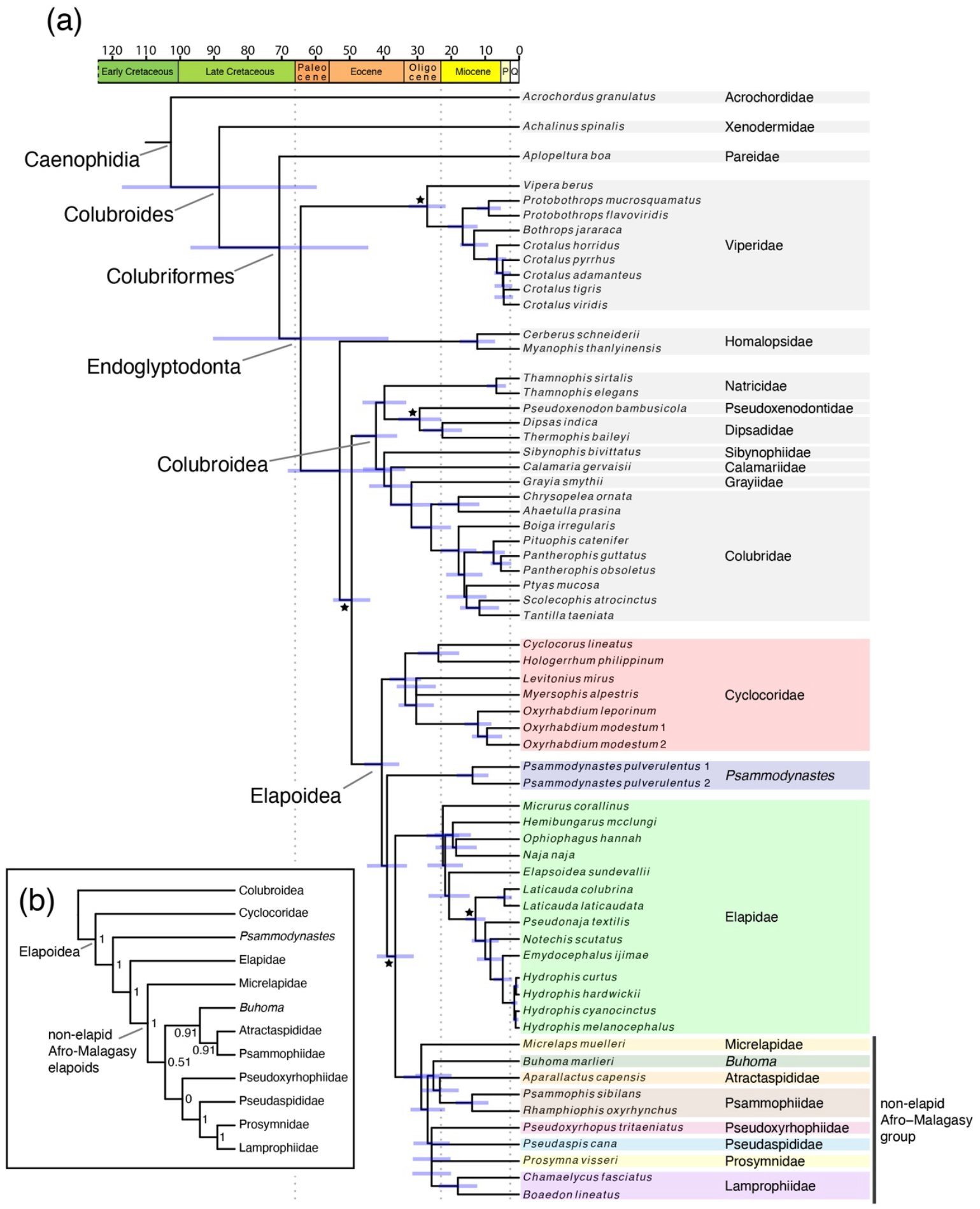
Coalescent-based species tree (a) with divergence times estimated from maximum likelihood node dating and time scale in millions of years, and cladogram (b) detailing topology of the species tree (a) for lineages of Elapoidea. Black stars indicate fossil calibrated nodes; blue bars indicate divergence time confidence intervals; P = Pliocene Epoch; Q = Quaternary Period; numbers at internal nodes of (b) indicate local posterior probability (PP) support for monophyly and PP ≥ 0.95 was considered strong support. This figure was illustrated using R packages ‘ggtree’ 3.0.4 and ‘deeptime’ 0.2.2 [168,169].

### 3.4. Biogeography

Results of biogeographic analyses supported an Asian origin for both Colubroidea and Elapoidea under all biogeographic models (figure 4; electronic supplementary material, figures S3–S8). Statistical comparison of results supported the BAYAREALIKE+J model as the optimal biogeographic model (AIC = 551). Under this model (figure 4; electronic supplementary material, figure S3), our results supported multiple successful colonisations by ancestral colubroid or elapoid lineages from Asia into Africa (n = 15), the Americas (n = 6), Australasia (n = 4), or Europe (n =5); from Africa into Asia (n = 7), Europe (n = 6), Australasia (n = 1); from Europe into Asia (n = 1); from land habitats in Australasia to marine habitats (n = 2); and into Europe by ancestral colubroid lineages that were inferred as being widely distributed across Africa and Asia (n = 2). Possibly the result of our lower sampling of Old World versus New World ratsnakes (Coronellini), all models supported a single colonisation event from the Americas into Europe and Africa by an ancestor of *Coronella* (Colubridae), which we consider unlikely comparted to an alternative scenario involving colonisation by ancestral *Coronella* from Asia into Europe and Africa. This latter scenario is consistent with biogeographic results of an earlier study [88] that included species-level phylogenomic sampling for ratsnakes.

**Figure 4.**
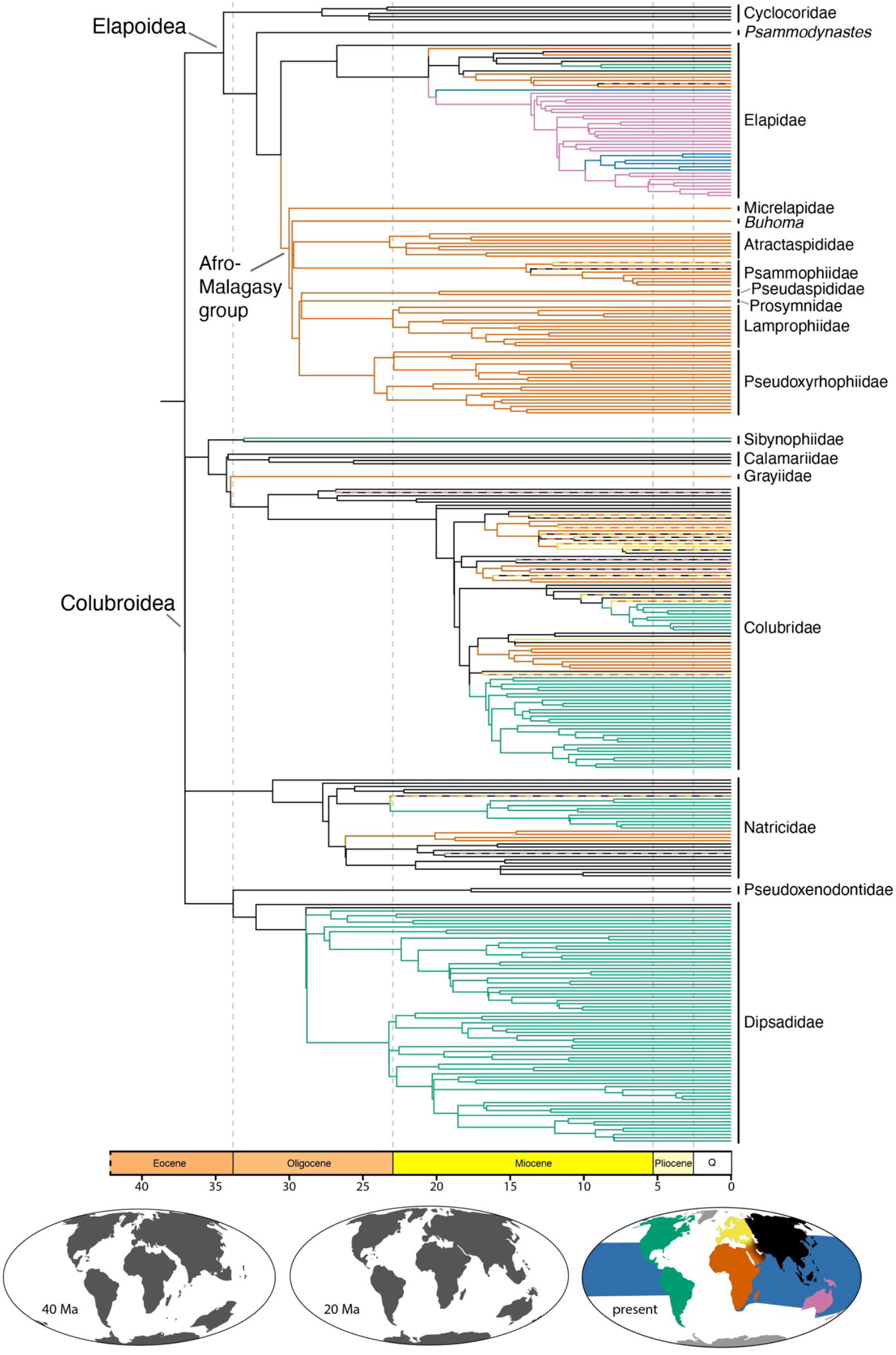
Phylogeny of Elapoidea and Colubroidea with genus-level sampling and estimates of historical geographic ranges. Branch colours correspond to colours in map and indicate maximum posterior probability estimate of geographic range of descendant node. Time scale is in millions of years. Paleomaps in this figure were illustrated using R packages ‘rgplates’ 0.2.0 and ‘chronosphere’ 0.4.1 [170,171].

Within Elapoidea, our results supported four ‘Out of Asia’ colonisation events into Africa, including: (1) by an ancestor of the Afro-Malagasy group approximately 24.4–37.5 Ma (95% CI), which spans the latest Eocene and most of the Oligocene; (2) by an ancestor of the lineage containing African cobras (genera *Aspidelaps, Hemachatus, Naja, Pseudohaje*, and *Walterinnesia*) approximately 12.5–23.9 Ma (early and middle Miocene); (3) by an ancestor of African Garter Snakes (genus *Elapsoidea*) as early as 25.6 Ma; and (4) by an ancestor of Mambas (genus *Dendroaspis*) as early as 18.9 Ma. After these lineages colonised and diversified within Africa, our results supported successful ‘Out of Africa’ colonisation of Europe by at least one sublineage in genus *Malpolon*, as early as 17.1 Ma, and separate recolonisations of Asia by *Psammophis* (<17.9 Ma) and *Naja* (<15.1 Ma).

Maximum likelihood ancestral geographic ranges estimated with biogeographic model DIVALIKE+J (AIC = 555; electronic supplementary material, figure S4) were identical to our estimates under the optimal model. Ranges estimated with model DEC+J (AIC = 1217; electronic supplementary material, figure S5) differed from ranges estimated under the optimal model by inferring an earlier colonisation of Australasia from Asia, which involved subsequent, additional colonisations from Australasia to Africa and back to Asia; and an earlier colonisation of Africa, which involved an additional colonisation back to Asia. Model DEC (AIC = 598; electronic supplementary material, figure S6) supported two colonisations of Australasia by ancestral elapoids in Asia (versus one colonisation under the optimal model) and two colonisations of marine environments from land habitats in Asia (versus from land habitats in Australasia under the optimal model). Furthermore, compared to our results under the optimal biogeographic model, our results under models BAYAREALIKE (AIC = 729; electronic supplementary material, figure S7) and DIVALIKE (AIC = 627; electronic supplementary material, figure S8) supported only one successful colonisation of Africa by an elapoid: 27.9– 38.6 Ma, from Asia, by an ancestor of the clade containing all elapoids except Cyclocoridae, which involved additional, subsequent colonisations of Asia from Africa.

## 4. Discussion

### 4.1. Overview

We inferred robust estimates of phylogenetic relationships and of geographic range evolution for the snake superfamily Elapoidea, a group distributed worldwide in tropical and subtropical areas that has undergone multiple rounds of rapid lineage diversification, and that includes many dangerously venomous species (e.g., Cobras, Coral Snakes, and Sea Snakes), as well as species that are harmless or not known to be medically significant to humans [89–91]. We generated, for the first time to date, a sub-genomic (sequence capture) dataset sampling all major elapoid lineages. Our phylogenetic and divergence time estimates (figures 2–3) confirm early, successive divergences of the two Asia-endemic lineages, Cyclocoridae and *Psammodynastes*, within Elapoidea. Our geographic range evolution inference (figure 4) is best interpreted as supporting an Asian origin for Elapoidea, with multiple faunal exchanges between Asia and either Africa, Australasia, the Americas, or Europe.

### 4.2. Biogeography

#### ‘Out of Africa’ hypothesis not supported for stem Elapoidea

Our phylogenetic and biogeographic results supported an Asian origin for each of superfamilies Colubroidea and Elapoidea (figure 4)—inconsistent with the ‘Out of Africa’ hypothesis previously suggested for Elapoidea [25] and supported in biographical literature for other vertebrate groups [30–37]. Fossil occurrences indicate an early elapoid presence in Africa: the oldest and only known Paleogene fossils that have been interpreted as representative of Elapoidea are from southern Tanzania, 25 ± 0.1 Ma, during the Late Oligocene [38,92]. As such, and given the variable and conflicting phylogenetic results from earlier Sanger-based DNA sequence data sets, the Kelly *et al*. [25] ‘Out of Africa’ hypothesis has remained plausible. In the most recent phylogenomic studies [22,23], however, an Asian origin was supported for Elapoidea (if not discussed), given that Cyclocoridae (endemic to the Philippines) and *Psammodynastes* (endemic to eastern Asia) were recovered as the earliest branching elapoid lineages, and *Calliophis* (endemic to Asia) has been estimated often as the earliest branching lineage within Elapidae [24,26,28,93]. Our results definitively supported an Asian origin for Elapoidea, rejecting the ‘Out of Africa’ hypothesis in support of the Asian origins of this group.

The absence of Paleogene elapoid fossils outside of Africa may reflect geographic and temporal sampling gaps and uncertainty in assignment of fossils to taxonomic groups and evolutionary lineages [94]. To avoid biases associated with *a priori* misassignment of fossils to nodes or lineages during phylogenetic inference, total evidence phylogenetic dating methods for integrated analysis of molecular and morphological data have been used to jointly estimate divergence times and evolutionary relationships for extant and extinct species of other vertebrate groups [95,96], and application of these methods in future biogeographic studies of Elapoidea may provide further insight on phylogenetic relationships, divergence times, and geographic range evolution in this group. Regardless, we consider our phylogenetic and biogeographic results to be strong evidence for an Asian origin for both Colubroidea and Elapoidea, a scenario that necessitates the related interpretation that numerous faunal exchanges explain the current presence of elapoid species throughout Asia, Africa, Australasia, the Americas, and Europe.

#### Faunal exchanges between Asia and the Americas

The modern terrestrial vertebrate fauna of the Americas includes ancient lineages that survived breakup of Laurasia or Gondwana and more recent arrivals that colonised the region by dispersal through land bridge corridors and/or over marine channels, or possibly even across open ocean [14,24,28,97–102]. Fossil evidence and molecular phylogenetic studies have supported the Bering Land Bridge (BLB) and former North Atlantic land bridges as major dispersal corridors between Asia or Europe and the Americas, respectively [97,103–105], but whether or not these land connections were also dispersal corridors for snakes is still unclear, given that fossils of snakes are not yet known from areas coinciding with locations of former land bridges [94,106,107]. Previous phylogenetic results [97,99,108–110] and our time-calibrated phylogenomic and biogeographical range evolution results (figures 3–4) lend support to the interpretation of multiple colonisations from Asia, including by at least one elapoid lineage and multiple colubroid lineages. In all instances inferred here, phylogenetic relationships now firmly point to their origins in Asia, followed by separate colonisations of western North America, possibly via a BLB-facilitated overland dispersal conduit [108].

Land connections between western Europe and eastern North America are thought to have been submerged >50 Ma [103]—prior to hypothesized colonisations by elapoid and colubroid lineages studied here (figure 3*a*)—whereas Beringia, during periods such as the Miocene Climatic Optimum (15–17 Ma), harbored lineages usually associated with warm climates [111–113], which may have contained habitats suitable for ectothermic vertebrates. Our phylogenetic and geographic range evolution results are consistent with historical dispersals across a Beringia land corridor, but snake fossils are net yet known from this region [94,106,107] and as such, our results cannot be interpreted as evidence against oceanic dispersal colonisation [114].

#### Faunal exchanges between Asia and Australasia

Hypothesized mechanisms for biotic exchange between Australasia and Asia have necessarily considered trans-oceanic dispersal, either directly or by island-hopping along intervening island chains [115,116]. Alternatively, paleotransport, whereby lineages are thought to have rafted on drifting islands or continental fragments over geological time scales [117], may explain Asia–Australasia biotic disjunctions, as may a combination of these processes [115,118–121]. The highly dynamic, complex tectonic landscape encompassing the region between Asia and Australasia makes paleogeographical modelling difficult [122,123], which, combined with statistical uncertainty in all estimates of lineage ages and faunal exchange times, makes it difficult to distinguish between alternative faunal exchange mechanisms between these regions. Nevertheless, our results strongly support immigration to Australasia at least once by an elapoid sublineage from Asia, and consistent with earlier studies, by multiple colubroid sublineages [36,124,125]. Our biogeographic range evolution results strongly supported successful colonisation of Australasia from Asia by an ancestor of the group containing all extant Australasian elapids and sea snakes—a group classically recognized as an adaptive radiation [126,127]. Although our biogeographical range evolution results supporting two origins for extant sea snakes from ancestrally terrestrial lineages (Hydrophiinae and Laticaudinae) are consistent with the prevailing hypothesis regarding their origins, future studies should investigate whether it is appropriate to use traditional biogeographic models (e.g., DEC) for cases such as land-to-marine transitions, where colonisation likely requires adaptive evolution [128–130]. Following ancestral adaptive colonisations from land to sea, both sea snake lineages further diversified in environments of the Indian and western Pacific Oceans [131,132].

#### Faunal exchanges between Africa and Eurasia

Hypothesized mechanisms for Cenozoic faunal exchange between Africa and Asia or Europe have included trans-oceanic dispersal across the Tethyan Seaway, Mediterranean Sea, or Indian Ocean, or dispersal across either the *Gomphotherium* land bridge resulting from Africa-Eurasia collision approximately 20 Ma [133] or an ephemeral land bridge associated with the Messinian Salinity Crisis [134–136]. Previous molecular phylogenetic studies and fossil evidence have supported a trans-Tethyan dispersal mechanism for some lizards [137], tortoises [138,139], and multiple mammal lineages [140,141]. Our biogeographic range evolution results were consistent with trans-Tethyan dispersal as the mechanism of faunal exchange only for the non-elapid Afro-Malagasy group (figure 4).

Das *et al*. [22] inferred a middle-to-late Eocene origin for the non-elapid Afro-Malagasy group, consistent with a trans-Tethys dispersal history, and interpreted rapid diversification in this group as indicative of increased ecological opportunity following the Cretaceous-Paleogene (K-Pg) mass extinction event. In contrast, our results (figures 3–4) and those of some earlier studies [28,142] supported more recent (late Eocene to Oligocene; 24–37 Ma) colonisation and rapid diversification, and, therefore, call into question previous support [22] for the K-Pg extinction-associated ecological opportunity hypothesis. We cannot entirely reject the hypothesized association between K-Pg extinction and rapid diversification, however, and Africa is thought to have had a depauperate Paleogene vertebrate fauna until Africa-Eurasia collision near the end of the Oligocene [133]. Additional field sampling and analyses of fossils are needed to characterize Paleogene snake communities in Africa to better assess the long-term impacts of K-Pg extinction on species diversity, ecological opportunity, and rapid diversification in lineages that subsequently colonised the continent [143,144].

Wüster *et al*. [145] inferred a mid-Eocene origin for the African cobra lineage encompassing *Aspidelaps, Hemachatus, Naja, Pseudohaje*, and *Walterinnesia* consistent with a trans-Tethyan dispersal mechanism for their colonisation, whereas our estimates support a more recent (mid-to-late Miocene; 12.5–23 Ma) arrival for cobras in Africa (figure 4), encompassing the time period when Africa and Eurasia collided and the *Gomphotherium* land bridge formed [146]. This land connection has been implicated as a major corridor for faunal exchange between Africa and Eurasia, which Rage & Gheerbrant [133] referred to as the Great Old World Interchange, and may have also facilitated expansions into Africa by ancestral populations of African Garter Snakes (*Elapsoidea*) and mambas (*Dendroaspis*). After elapoid lineages successfully colonised Africa and diversified, our phylogenetic and biogeographic results (figures 3–4) supported multiple ‘Out of Africa’ colonisations of Asia or Europe by ancestral *Psammophis, Malpolon*, and *Naja* sublineages.

Previous phylogenetic and phylogeographic studies have supported colonisation of Asia at least twice by lineages of *Psammophis* (Psammophiidae), first by an ancestor of the clade minimally containing *Psammphis condanarus, P. indochinensis, P. lineolatus*, and *P. turpanensis*, and probably also containing *P. leithii* and *P. longifrons* [147,148]; and more recently by either an ancestor of *Psammophis schokari* or by an ancestor of the Asian sublineage within this species [149]. Whether or not *Psammophis leithii* and *P. longifrons* represent additional invasions of the continent must be tested with genetic data. Kurniawan *et al*. [148] hypothesized that the earlier of these *Psammophis* immigration events occurred by dispersal across the *Gomphotherium* land bridge, and our biogeographic and divergence time results were consistent with this hypothesis (figure 4). The importance of *Psammophis* fossils from the Late Miocene of Spain [150]—where the genus does not presently occur—in understanding when and how ancestors of extant *Psammophis* lineages arrived in Asia is not clear.

Our estimates of divergence times and biogeographic range evolution results were also consistent with colonisation of Europe by an ancestor or sublineage of *Malpolon* as early as the mid-to-late Miocene (5.5–12.3 Ma). However, this early estimate is likely an artifact of our sampling only one lineage for the genus. Previous phylogeographic studies that had dense geographic genetic sampling supported multiple recent arrivals of the genus into Europe, including at least once by a sublineage of *Malpolon monspessulanus*, likely across the Strait of Gibraltar during the Pliocene or Pleistocene [151], and separately during the Pleistocene by a sublineage of *M. insignatus* along a route east of the Mediterranean Sea [152]. Additional biotic exchanges between Africa and Eurasia have been supported for numerous vertebrate lineages during the late Miocene, and may have been facilitated by the presence of volcanic island arcs [134] and ephemeral land bridges spanning the Mediterranean during the Messinian Salinity Crisis [135,136]. The propensity of snakes for over water dispersal, as suggested by their current presence on many oceanic islands [7,77,78,81,153–156], suggests that additional oceanic dispersal-facilitated colonisations may have occurred over deep time by lineages that have since gone extinct.

### 4.3. Phylogenetic relationships

Our robust estimate of evolutionary relationships (figure 2) is the first to include all major lineages of Elapoidea (earlier studies did not include *Buhoma, Micrelaps*, and *Psammodynastes* together in a phylogenomic dataset) and is congruent with phylogenetic results of several recent studies that sampled hundreds or thousands of loci [22,23,27]. Our phylogenetic results provide additional, strong statistical support for monophyly of the “Afro-Malagasy group” comprising 330 species with a diversity of phenotypes and ecologies (figures 2–3).

On the basis of their phylogenetic results and interpretation of skeletal synapomorphies uniting *Micrelaps* and *Brachyophis* (no molecular data yet exists for the latter genus), Das *et al*. [22] proposed a new taxonomic family, Micrelapidae, within which they included *Micrelaps* and *Brachyophis*. Our phylogenetic results, which inferred *Micrelaps* as the earliest diverging lineage within the Afro-Malagasy group, were congruent with results of Das *et al*. [22], and, therefore, we also recognize Micrelapidae as a family-level lineage. We are preparing a manuscript to formally establish family-rank taxonomic groups for *Buhoma* and *Psammodynastes*, considering that our phylogenetic results strongly supported these taxa as lineages that are genetically distinct from each other and from other elapoid lineages.

Resolving elapoid evolutionary relationships has been notoriously difficult due to rapid diversification in the group [22,25]—characterized in our inferred time trees by the presence of many short internal branches early in the history of the Afro-Malagasy group (figures 3–4). Rapid diversification is often associated with high genealogical discordance resulting from increased ILS and/or historical hybridization, prevalent in the evolutionary history of many vertebrate groups [157–165]. Results of our ILS and hybridization analyses (electronic supplementary material, figure S2 and table S5) were consistent with ILS-driven genealogical discordance and did not support historical hybridization events between major elapoid lineages. Although hybridization has been hypothesized to contribute to rapid diversification [162], our inferred lack of support for hybridization is inconsistent with this hypothesis for Elapoidea, at least for the deeper elapoid lineages sampled herein. Erosion of genomic signatures of hybridization, however, can make detection of ancient hybridization events difficult [166].

### 4.4. Future directions

Future investigations of elapoid evolutionary relationships, divergence times, and ancestral geographic ranges should use phylogenomic datasets that include more species to assess prevalence of historical hybridization, because hybridization can impact evolutionary relationships inferred using tree-based phylogenetic methods and downstream inferences from results of ancestral biogeographic reconstruction analyses. To assess whether extinct lineages (represented in the fossil record) can be included in robust phylogenetic estimates, integrative phylogenetic analysis using molecular and morphological datasets should be explored. Considering that most extinct snake species are known only from isolated vertebra, investigations should assess whether vertebral elements contain enough phylogenetic signal for robust phylogenetic inference, as previously suggested by Smith & Georgalis [94].

Our time calibrated phylogenetic estimates and ancestral geographic range estimates will contribute to comprehensive investigations of historical environmental change. Such studies, if focused within hypothesized faunal exchange corridors, such as Beringia, would facilitate testing of “shared process” hypotheses related to historical biotic exchange. Characterization of paleoenvironments, increased sampling in future phylogenetic datasets of Elapoidea, and construction of large phenotypic and ecological datasets would enable tests of hypotheses related to drivers of lineage diversification.

## 5. Conclusions

Our phylogenetic results provided strong statistical support for relationships that historically have been difficult to resolve, while confirming relationships recovered in recent phylogenomic studies, highlighting the usefulness of generating phylogenomic datasets by targeting loci expected to contain high phylogenetic signal. Moreover, our results demonstrate the advantage of using phylogenomic datasets, compared to using datasets comprising a small number of easy-to-sequence “phylogenetic marker” loci, when inferring relationships among lineages that rapidly diversified.

Biogeographic analyses using our improved phylogenetic estimate of relationships within Elapoidea unanimously supported an Asian origin for this diverse clade and strongly rejected the ‘Out of Africa’ scenario suggested in earlier phylogenetic studies of the group. This rejection of the ‘Out of Africa’ history in favor of the ‘Out of Asia’ history for ancestral Elapoidea largely reflects our improved estimate of evolutionary relationships in the group. Our novel biogeographic results highlight how prior phylogenetic assumptions can strongly impact ancestral reconstruction analyses. As large phylogenomic datasets become available for a broader diversity of biota, our ability to test major evolutionary and biogeographic hypotheses will drastically improve.

## Acknowledgments

We thank the Philippine Department of Environment and Natural Resources (DENR) and the Biodiversity Management Bureau (BMB) for facilitating collecting and export permits necessary for components of the present study. For loans of genetic samples we thank Aaron Bauer (Villanova University), Eli Greenbaum (University of Texas, El Paso), and Shai Meiri (Steinhardt Museum of Natural History, Tel Aviv University). We thank Jennifer Raff, Kirsten Jensen, and Rob Moyle for their input on an earlier version of this work.

## Ethics Statement

Tissue samples were obtained using methods approved by the University of Kansas Institutional Animal Care and Use Committee.

## Data Accessibility

Raw sequence reads of targeted sequence capture data were accessioned in the NCBI Sequence Read Archive with BioProject PRJNA926108; sample and GenBank accession IDs associated with novel sequence data are included in electronic supplementary material [167]. Sequence alignments, geographic data, and phylogenetic estimates are available in Open Science Framework project krhx3.

## Declaration of AI use

We have not used AI-assisted technologies in creating this article.

## Authors’ Contributions

JLW: conceptualization, field work, formal analysis, funding acquisition, writing— original and revised drafts; FTB: conceptualization, formal analysis, writing—review and editing; SD: conceptualization, writing—review and editing; RMB: conceptualization, field work, funding acquisition, writing—review and editing. All authors read and gave final approval for publication of this manuscript.

