## Supplemental Materials for "Novel phylogenomic inference and ‘Out of Asia’ biogeography of cobras, coral snakes, and their allies"

**Electronic Supplementary Material for "Novel phylogenomic inference and 'Out of Asia' biogeography of cobras, coral snakes, and their allies"**

**Tissues.** Tissues were collected by J. Alesio, K. E. Allen, V. Arias, R. M. Brown, W. and J. Bulalacao, L. Canseco, A. C. Diesmos, E. Ducasse, W. E. Duellman, J. B. Fernandez, E. B. Greenbaum, C. Hayden, J. Guayasamin, B. Gurubat, D. S. McLeod, J. Porras, S. Pouy, C. Raxworthy, R. Reyes, D. Roldán-Piña, E. L. Rico, M. B. Sanguila, C. D. Siler, J. E. Simmons, W. P. Tapondjou N., I. Tigulu, S. L. Travers, L. J. Welton, E. and Z. Wickham, and J. Zafe during field expeditions in Cameroon (2015, 2018), China (2006, 2007), El Salvador (2000), Ghana (1998), Paraguay (1996), Peru (1990), Philippines (2000, 2001, 2005, 2007, 2008, 2009, 2011, 2013, 2014, 2017), Solomon Islands (2016), and Thailand (2005), and deposited in the University of Kansas Biodiversity Institute, or as loans from tissue collections at University of Texas El Paso and Villanova University (Villanova, PA).

**Selecting loci to target for sequence capture.** Using a probe-based sequence capture approach, we targeted rapidly evolving exons (REEs), ultraconservative elements (UCEs), ddRAD-like loci (inferred *in-silico*), and protein-coding regions of previously identified major histo-compatability (MHC), vision, and scalation-associated genes. To identify REEs, we used the R package REEs v1.9 [1] and 11 previously published squamate genome assemblies (https://github.com/JeffWeinell/SnakeCap). To select ddRAD-like loci, we used the REEs function "proposeLoci.ddRADlike" to locate 900–1000bp regions in the genome of *Thermophis baileyi* that begin and end with the *EcoR1* and *SbfI* restriction enzyme site recognition sequences (CCTGCAGG and GAATTC), respectively. To select vision loci, we used BLASTn from BLAST+ v2.9 [2] to search within sampled snake genomes for matches to probe sequences used by [3] for vision-associated genes in *Anolis*, *Columba*, *Gallus*, *Pelodiscus*, *Sceloporus*, or *Python*, and our probes corresponded to the genomic hit sequences of best matches. To select scalation loci, we used tBLASTn from BLAST+ to search for *Ophiophagus* and *Python* Epidermal Differentiation Complex protein sequences previously identified by [4] in genomes of *T. sirtalis*, *P. mucrosquamatus*, and *C. horridus*. Furthermore, we included a subset of the UCEs of *Micrurus fulvius* from the study by [5], and 27 sequence regions corresponding to annotated major histocompatibility complex genes from the genome of *Thamnophis sirtalis*. Arbor Sciences synthesized 120nt RNA probes for our target sequences with 2x tiling and ultrastringent filtering, as a custom MyBaits 20,000 RNA-probes kit, and conducted library preparation prior to sequencing by NovoGene on an Illumina HiSeq X sequencing (dual-index, paired-end, 150 bp inserts). The optimized probe set included 20,020 probes for 3128 loci (1,517,011 nt) selected as targets: 1652 REEs, 907 UCEs, 328 ddRAD-like, 27 MHC, 119 vision, and 95 scalation loci (Open Science Framework, project krhx3).

**Geographic transition zones.** We considered two geographic transitional zones when coding taxa as present or absent in our geographic range dataset used for biogeographic range evolution analyses. One transition zone comprises the area containing Anatolia, Armenian Highlands, and Iranian Plateau, where Africa, Asia, and Europe meet. A second transition zone comprises islands of Wallacea between Asia and Australasia (figure S1). Colubroid or elapoid genera are not endemic to either transition zone [6–11], and therefore when scoring genera as present or absent in regions, we ignored portions of their ranges intersecting transition zones. For example, genera occurring both within the Africa-Asia-Europe transition zone and in Africa outside of the transition zone were coded in our geographic range dataset as occurring only in “Africa”.

**Table S1.** Individuals sampled using targeted sequence capture (bold font) or from previously published assembled genomes. Institution and field series abbreviations: KU = University of Kansas; Steinhardt Museum of Natural History, Tel Aviv University; MCZ = Museum of Comparative Zoology; PNM = Philippine National Museum; UTEP = University of Texas, El Paso. For sequence capture data, raw reads are available in the NCBI Sequence Read Archive with BioProject ID PRJNA926108. Sequence alignments used for phylogenetic analyses are available on Open Science Framework, project krhx3. Data ID is either the NCBI BioSample accession number (SAMN numbers), NCBI genome assembly accession number (GCA numbers), or 'snake_7C' for the *Boa constrictor* genome assembly (GigaDB repository; project: http://dx.doi.org/10.5524/100060).

| **Species** | **Family** | **voucher ID** | **other sample ID** | **data ID** |
|---|---|---|---|---|
| ***Acrochordus granulatus*** | Acrochordidae | KU 302952 | CDS 681 | SAMN32850298 |
| ***Achalinus spinalis*** | Xenodermidae | KU 312258 | MCB 516 | SAMN32850299 |
| ***Cerberus schneiderii*** | Homalopsidae | KU 324534 | CDS 5109 | SAMN32850300 |
| ***Aplopeltura boa*** | Pareidae | KU 315147 | RMB 10058 | SAMN32850301 |
| ***Ahaetulla prasina preocularis*** | Colubridae | KU 323364 | RMB 12330 | SAMN32850302 |
| ***Calamaria gervaisii*** | Calamariidae | KU 327406 | ACD 1871 | SAMN32850303 |
| ***Boiga irregularis*** | Colubridae | KU 345021 | SLT 833 | SAMN32850304 |
| ***Scolecophis atrocinctus*** | Colubridae | KU 289804 | EBG 276 | SAMN32850305 |
| ***Tantilla taeniata*** | Colubridae | KU 289863 | EBG 393 | SAMN32850306 |
| ***Dipsas indica*** | Dipsadidae | KU 214857 | WED 59279 | SAMN32850307 |
| ***Grayia smythii*** | Grayiidae | KU 341872 | RMB 19318 | SAMN32850308 |
| ***Pseudoxenodon bambusicola*** | Pseudoxenodontidae | KU 312221 | KU-FS 734 | SAMN32850309 |
| ***Sibynophis bivittatus*** | Sibynophiidae | KU 309608 | RMB 7779 | SAMN32850310 |
| ***Elapsoidea sundevallii*** | Elapidae | MCZ R-190390 | MCZ A-27807 | SAMN32850311 |
| ***Micrurus corallinus*** | Elapidae | KU 289205 | JES 1814 | SAMN32850312 |
| ***Hemibungarus mcclungi*** | Elapidae | KU 303027 | CDS 1486 | SAMN32850313 |
| ***Buhoma marlieri*** | *incertae sedis* | UTEP 22598 | CFS 1528 | SAMN32850314 |
| ***Aparallactus capensis*** | Atractaspididae | MCZ R-193258 | MCZ A-28808 | SAMN32850315 |
| ***Cyclocorus lineatus lineatus*** | Cyclocoridae | KU 335808 | RMB 17698 | SAMN32850316 |
| ***Oxyrhabdium leporinum*** | Cyclocoridae | KU 346583 | RMB 23647 | SAMN32850318 |
| ***Oxyrhabdium modestum*** | Cyclocoridae | KU 310866 | CDS 3006 | SAMN32850319 |
| ***Oxyrhabdium modestum*** | Cyclocoridae | KU 338101 | RMB 19059 | SAMN32850317 |

| | | | | |
|---|---|---|---|---|
| ***Levitonius mirus*** | Cyclocoridae | PNM 9872 | KU 337269 | SAMN32850320 |
| ***Hologerrhum philippinum*** | Cyclocoridae | KU 330752 | CDS 5735 | SAMN32850321 |
| ***Myersophis alpestris*** | Cyclocoridae | KU 308684 | ELR 1149 | SAMN32850322 |
| ***Chamaelycus fasciatus*** | Lamprophiidae | KU 348632 | KAE 243 | SAMN32850323 |
| ***Boaedon lineatus*** | Lamprophiidae | no voucher | EBG 705 | SAMN32850324 |
| ***Prosymna visseri*** | Prosymnidae | MCZ R-190223 | AMB 8075 | SAMN32850325 |
| ***Psammophis sibilans*** | Psammophiidae | KU 290424 | RAX 2053 | SAMN32850326 |
| ***Pseudaspis cana*** | Pseudaspididae | no voucher | MCZ A-27068 | SAMN32850327 |
| ***Pseudoxyrhopus tritaeniatus*** | Pseudoxyrhophiidae | KU 340969 | SML 029 | SAMN32850328 |
| ***Rhamphiophis oxyrhynchus*** | Rhamphiophiidae | KU 290451 | RAX 2104 | SAMN32850329 |
| ***Psammodynastes pulverulentus*** | *incertae sedis* | KU 329686 | RMB 14526 | SAMN32850330 |
| ***Psammodynastes pulverulentus*** | *incertae sedis* | KU 328547 | DSM 1384 | SAMN32850331 |
| ***Micrelaps muelleri*** | Micrelapidae | SMNH TAU R-16738 | TAU R-16738 | SAMN32850332 |
| *Boa constrictor* | Boidae | — | ERS218597 | snake_7C |
| *Chrysopelea ornata* | Colubridae | — | — | GCA_018340635.1 |
| *Pantherophis guttatus* | Colubridae | — | — | GCA_019457695.1 |
| *Pantherophis obsoletus* | Colubridae | — | — | GCA_012654085.1 |
| *Pituophis catenifer* | Colubridae | — | — | GCA_019677565.1 |
| *Ptyas mucosa* | Colubridae | — | — | GCA_012654045.1 |
| *Thamnophis elegans* | Natricidae | — | — | GCF_009769535.1 |
| *Thamnophis sirtalis* | Natricidae | — | — | GCA_001077635.2 |
| *Thermophis baileyi* | Natricidae | — | — | GCA_003457575.1 |
| *Emydocephalus ijimae* | Elapidae | — | — | GCA_004319985.1 |
| *Ophiophagus hannah* | Elapidae | — | — | GCA_000516915.1 |
| *Hydrophis curtus* | Elapidae | — | — | GCA_019472885.1 |
| *Hydrophis cyanocinctus* | Elapidae | — | — | GCA_019473425.1 |
| *Hydrophis hardwickii* | Elapidae | — | — | GCA_004023765.1 |
| *Hydrophis melanocephalus* | Elapidae | — | — | GCA_004320005.1 |
| *Laticauda colubrina* | Elapidae | — | — | GCA_015471245.1 |

| | | | | |
|---|---|---|---|---|
| *Laticauda laticaudata* | Elapidae | — | — | GCA_004320025.1 |
| *Naja naja* | Elapidae | — | — | GCA_009733165.1 |
| *Notechis scutatus* | Elapidae | — | — | GCF_900518725.1 |
| *Pseudonaja textilis* | Elapidae | — | — | GCF_900518735.1 |
| *Myanophis thanlyinensis* | Homalopsidae | — | — | GCA_017656035.1 |
| *Python molurus* | Pythonidae | — | — | GCA_000186305.2 |
| *Bothrops jararaca* | Viperidae | — | — | GCA_018340635.1 |
| *Crotalus adamanteus* | Viperidae | — | — | GCA_018446365.1 |
| *Crotalus horridus* | Viperidae | — | — | GCA_001625485.1 |
| *Crotalus pyrrhus* | Viperidae | — | — | GCA_000737285.1 |
| *Crotalus tigris* | Viperidae | — | — | GCA_016545835.1 |
| *Crotalus viridis* | Viperidae | — | — | GCA_003400415.2 |
| *Protobothrops flavoviridis* | Viperidae | — | — | GCA_003402635.1 |
| *Protobothrops mucrosquamatus* | Viperidae | — | — | GCA_001527695.3 |
| *Vipera berus* | Viperidae | — | — | GCA_000800605.1 |

**Table S2.** Individuals with novel Sanger sequence data. Sequence alignment available on Open Science Framework, project krhx3.

| taxon | voucher | field tag ID | GenBank (CYTB) | GenBank (CMOS) |
|---|---|---|---|---|
| *Acrochordus granulatus* | KU 302951 | CDS 357 | OQ290841 | OQ290863 |
| *Aplopeltura boa* | KU 319950 | ACD 3924 | OQ290842 | OQ290864 |
| *Boiga cynodon* | KU 344107 | RMB 21632 | OQ290843 | OQ290865 |
| *Calliophis bilineatus* | KU 309511 | RMB 7884 | OQ290844 | OQ290866 |
| *Cerberus schneiderii* | KU 302983 | CDS 309 | OQ290845 | OQ290867 |
| *Chrysopelea paradisi* | KU 310165 | RMB 8473 | OQ290846 | OQ290868 |
| *Coelognathus erythrurus* | KU 303024 | CDS 1126 | OQ290847 | OQ290869 |
| *Cyclocorus nuchalis* | KU 334469 | RMB 16039 | OQ290848 | OQ290870 |
| *Gonyosoma oxycephalum* | KU 304104 | RMB 5388 | OQ290849 | OQ290871 |
| *Hemibungarus gemianulis* | — | CMNH H794 | OQ290850 | OQ290872 |
| *Hologerrhum philippinum* | KU 328837 | RMB 13628 | OQ290851 | OQ290873 |
| *Laticauda colubrina* | KU 341149 | SLT 197 | OQ290852 | — |
| *Laticauda colubrina* | KU 351734 | NDR 145 | OQ290853 | OQ290874 |
| *Naja samarensis* | KU 320521 | CDS 3662 | OQ290854 | OQ290875 |
| *Opisthotropis typica* | KU 327424 | RMB 3111 | OQ290855 | OQ290876 |
| *Oxyrhabdium leporinum* | KU 306307 | CDS 1769 | OQ290856 | OQ290877 |
| *Pseudorabdion taylori* | KU 327215 | ACD 5357 | OQ290857 | OQ290878 |
| *Sibynophis geminatus* | no voucher | ELR 198 | OQ290858 | OQ290879 |
| *Stegonotus muelleri* | KU 328844 | RMB 8541 | OQ290859 | OQ290880 |
| *Trimeresurus flavomaculatus* | KU 310865 | CDS 2895 | OQ290860 | OQ290881 |
| *Tropidolaemus subannulatus* | KU 334489 | RMB 16352 | OQ290861 | OQ290882 |
| *Tropidonophis spilogaster* | KU 329682 | RMB 14471 | OQ290862 | OQ290883 |

**Table S3.** Polymerase chain reaction and sequencing primers used in this study. Novel CYTB primers (72.15 and 987R.20) were designed using Primer3 v0.4.0 [12].

| locus | primer name | sequence | direction | source |
|---|---|---|---|---|
| CYTB | L14910 | GACCTGTGATMTGAAAACCAYCGTTGT | forward | [13] |
| CYTB | L14919 | AACCACCGTTGTTATTCAACT | forward | [13] |
| CYTB | H16064 | CTTTGGTTTACAAGAACAATGCTTTA | reverse | [13] |
| CYTB | H15720 | TGGTGTTGTGTAATTTGGTCT | reverse | [13] |
| CYTB | 72F.18 | AGCAATYCACTAYACAGC | forward | This study |
| CYTB | 337F.23 | TTCTGAGCAGCAACAGTAATCAC | reverse | [14] |
| CYTB | 732R.21 | YTCTGGTTTAATGTGTTGKGG | forward | [14] |
| CYTB | 987R.20 | AATGGAGGTTGTTTGACCAA | reverse | This study |
| C-MOS | S77 | CATGGACTGGGATCAGTTATG | forward | [15] |
| C-MOS | S78 | CCTTGGGTGTGATTTTCTCACCT | reverse | [15] |

**Table S4.** Fossils used to calibrate node ages for divergence time estimation.

| **Fossil description** | **Calibrated node** | **Fossil age (Ma)** |
|---|---|---|
| *Vipera* cf. *V. antiqua* Szyndlar & Böhme, 1993 | stem-Viperinae | 22.1 |
| Elapidae indet. McCartney, Stevens, & O'Connor, 2014 | stem-Elapidae | 24.9 |
| *Paleoheterodon tiheni* Holman, 1964 | stem-Dipsadidae | 12.5 |
| Colubridae indet. Smith, 2013 | stem-Colubroidea | 35.2 |
| *Incongruelaps iteratus* Scanlon, Lee, & Archer, & 2003 | stem-Oxyuraninae | 10 |

**Table S5.** Number of reticulations, complexity, and maximum pseudolikelihood of estimated phylogenetic networks; complexity (C) considered equal to number of identifiable (= number of internal branches + number of reticulations). Best model selected using slope heuristics estimated with capushe R package shown in bold, and using data-driven slope estimation (DDSE) and dimension jump (Djump) algorithms; MPL= maximum pseudolikelihood; MPL networks had at 0–4 reticulations, although maximum number of reticulations allowed was 0–9; HMAX = maximum number of reticulations allowed during network optimization; HMPL = number of reticulations in MPL tree.

| HMAX | HMPL | C | MPL | Notes |
|---|---|---|---|---|
| 0 | 0 | 9 | 87.172 | Results: This was the model chosen by capushe as optimal, under the jump_model algorithm, when complexity was set to observed (rather than maximum) number of hybridizations. |
| 1 | 1 | 10 | 72.340 | *Buhoma* + *Elaposidea* = stem *Psammophis* + *Aparallactus* |
| 2 | 2 | 11 | 65.567 | rerooting not possible |
| 3 | 3 | 12 | 52.966 | (*Elapsoidea* + *Aparallactus* = *Psammophis*); (*Boaedon* contribution to ancestral *Pseudoxyrhopus*); (*Psammodynastes* contribution to ancestral Africa-Malagasy lineage) Results: capushe selected this as the optimal model under both the DDSE_model and jump_model algorithms, when complexity was set to the maximum number of reticulations rather than to the observed number. |
| 4 | 4 | 13 | 54.421 | Re-rooting not possible on *Pseudoxenodon*, so rooted on *Hologerrhum*. *Hologerrhum* + *Elapsoidea* = *Pseudoxenodon*; *Buhoma* + *Psammophis* = *Aparallactus*; *Boaedon* + *Pseudaspis* = *Prosymna*; *Boaedon* + *Pseudaspis* + *Prosymna* stem contributed to *Buhoma* + *Psammophis* + *Aparallactus* stem. |
| 5 | **3*** | 12 | 52.943 | Not possible to reroot |
| 6 | **4*** | 13 | 54.171 | Not possible to reroot; Results: This was the model chosen by capushe as optimal, under the DDSE_model algorithm, when complexity was set to observed (rather than maximum) number of hybridizations. |
| 7 | 3* | 12 | 50.547 | *Elapsoidea* + *Aparallactus* = *Psammophis*; *Prosymna* + *Pseudoxyrhopus* = *Boaedon*; *Hologerrhum* + stem Africa-Malagasy lineage = *Psammodynastes* |
| 8 | 3* | 12 | 53.344 | *Psammophis* + stem Africa-Malagasy lineage = *Elapsoidea*; *Psammodynastes* contributed to *Hologerrhum*; *Prosymna* + *Pseudoxyrhopus* = *Boaedon* |
| 9 | 3* | 12 | 50.812 | *Hologerrhum* + stem-Africa Malagasy lineage = *Psammodynastes*; *Prosymna* + *Pseudoxyrhopus* = *Boaedon*; *Elapsoidea* + *Aparallactus* = *Psammophis*. |
| 10 | 3* | 12 | 50.658 | identical to max H = 9 network |

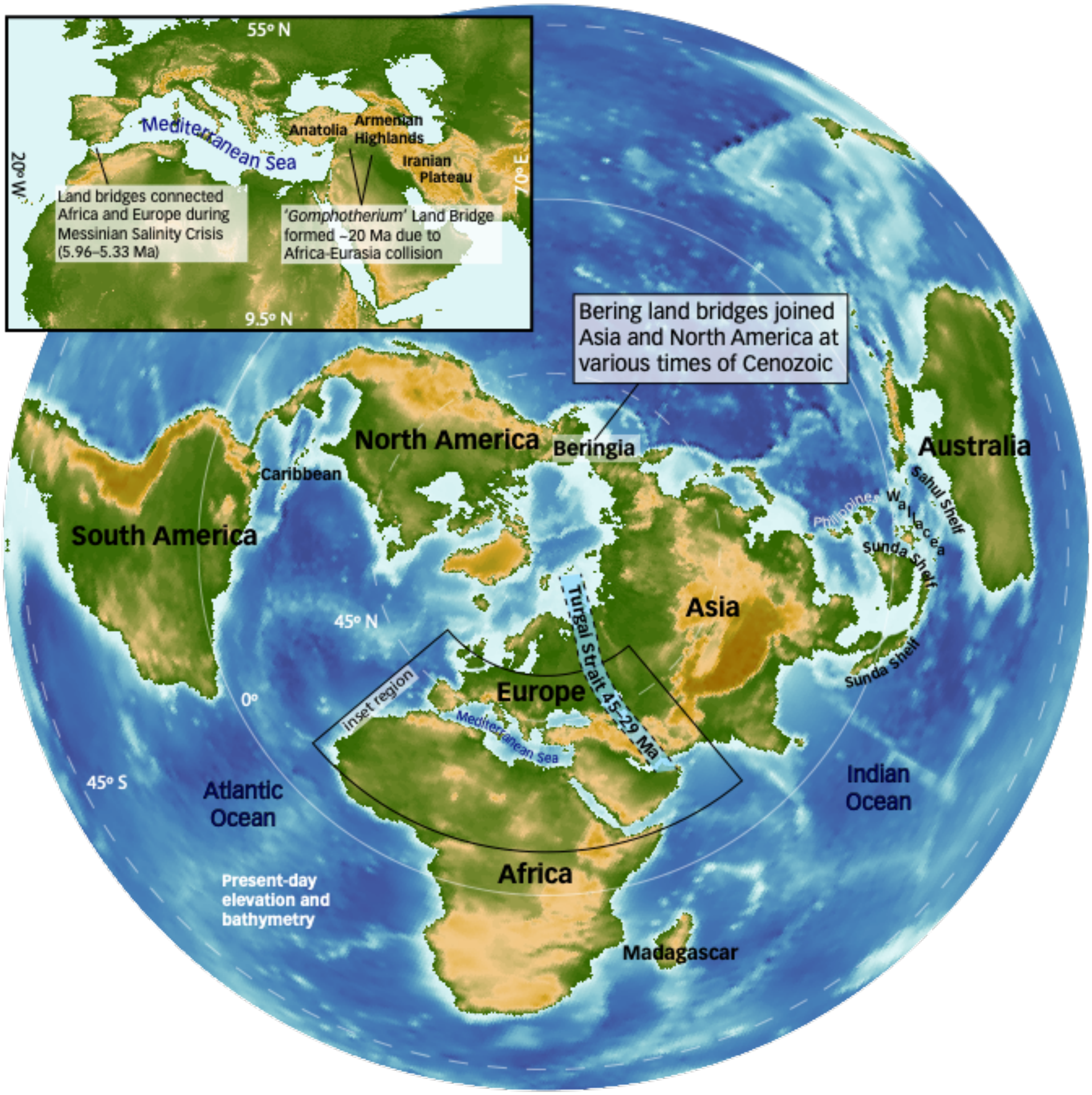


**Figure S1.** Places mentioned in this article. Colors indicate elevation above sea level (increasing from green to yellow to brown) and marine depth (blues; darker = deeper). Paleogeographical features hypothesized as facilitating or limiting historical faunal exchanges are described and indicated in the context of modern geography.

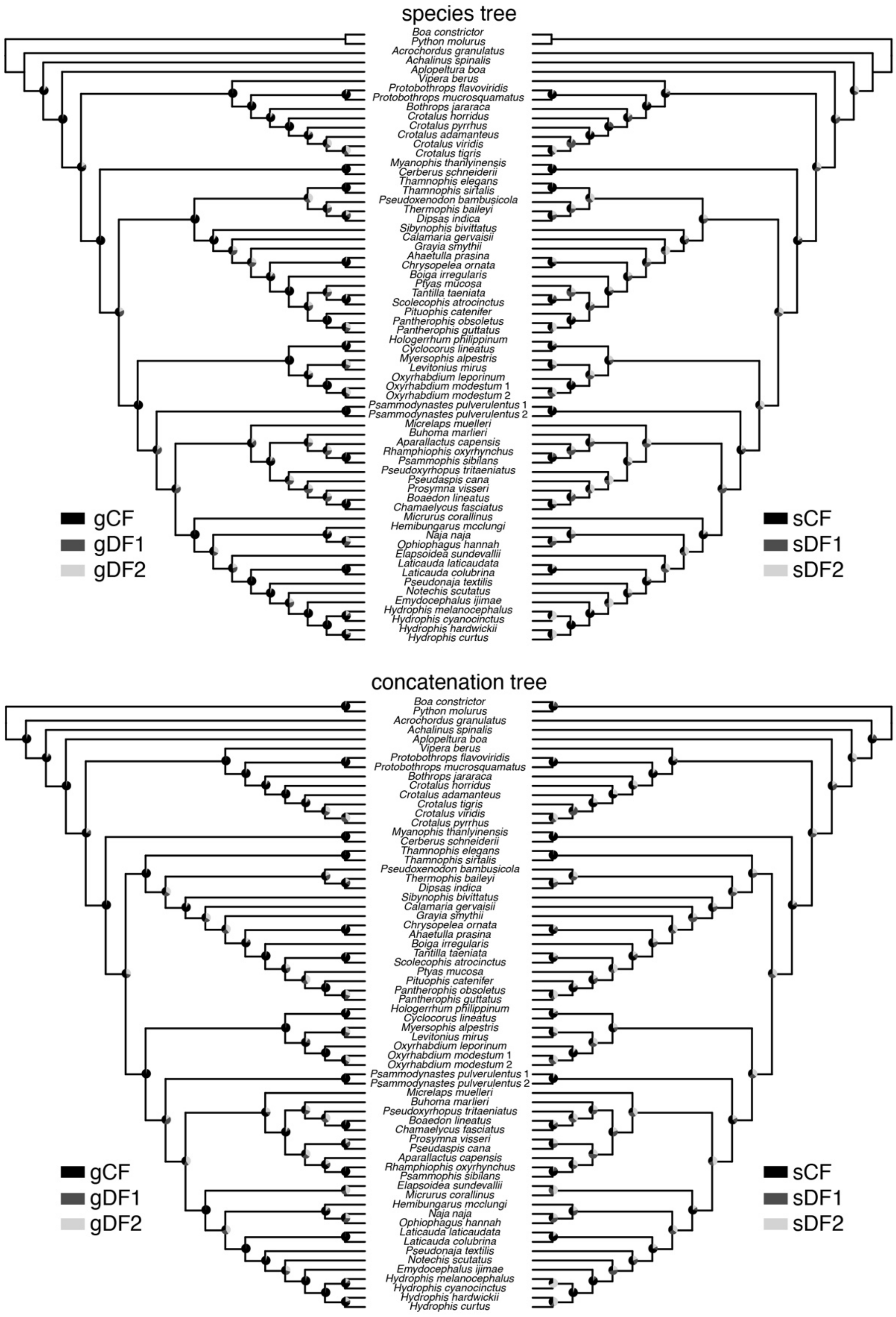


**Figure S2.** Species tree (top row) and concatenation tree (bottom row) with pie charts at nodes indicating gene (left) and site (right) concordance and discordance factors; gCF = gene concordance factor, gDF1 = gene discordance factor 1, gDF2 = gene discordance factor 2; sCF = site concordance factor, sDF1 = site discordance factor 1, sDF2 = site discordance factor 2.

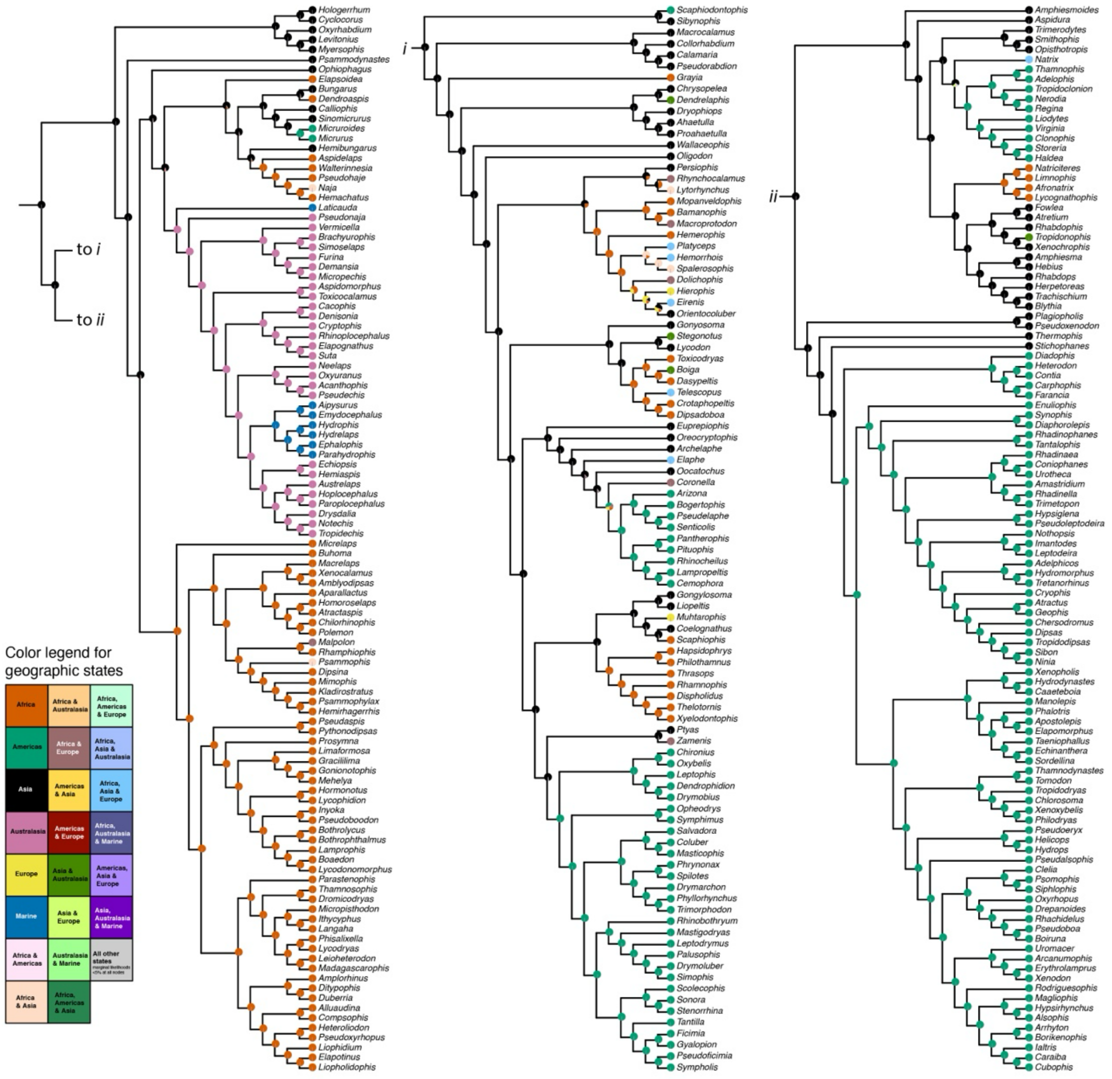


**Figure S3.** BioGeoBEARS geographic range evolution results of Elapoidea and Colubroidea under the biogeographic model “BAYAREALIKE+J”. Node pie charts depict marginal probability estimates of ancestral geographic states. Tip circles depict geographic states of extant lineages provided as input during analysis. Phylogeny used during analysis is visualized here as a cladogram.

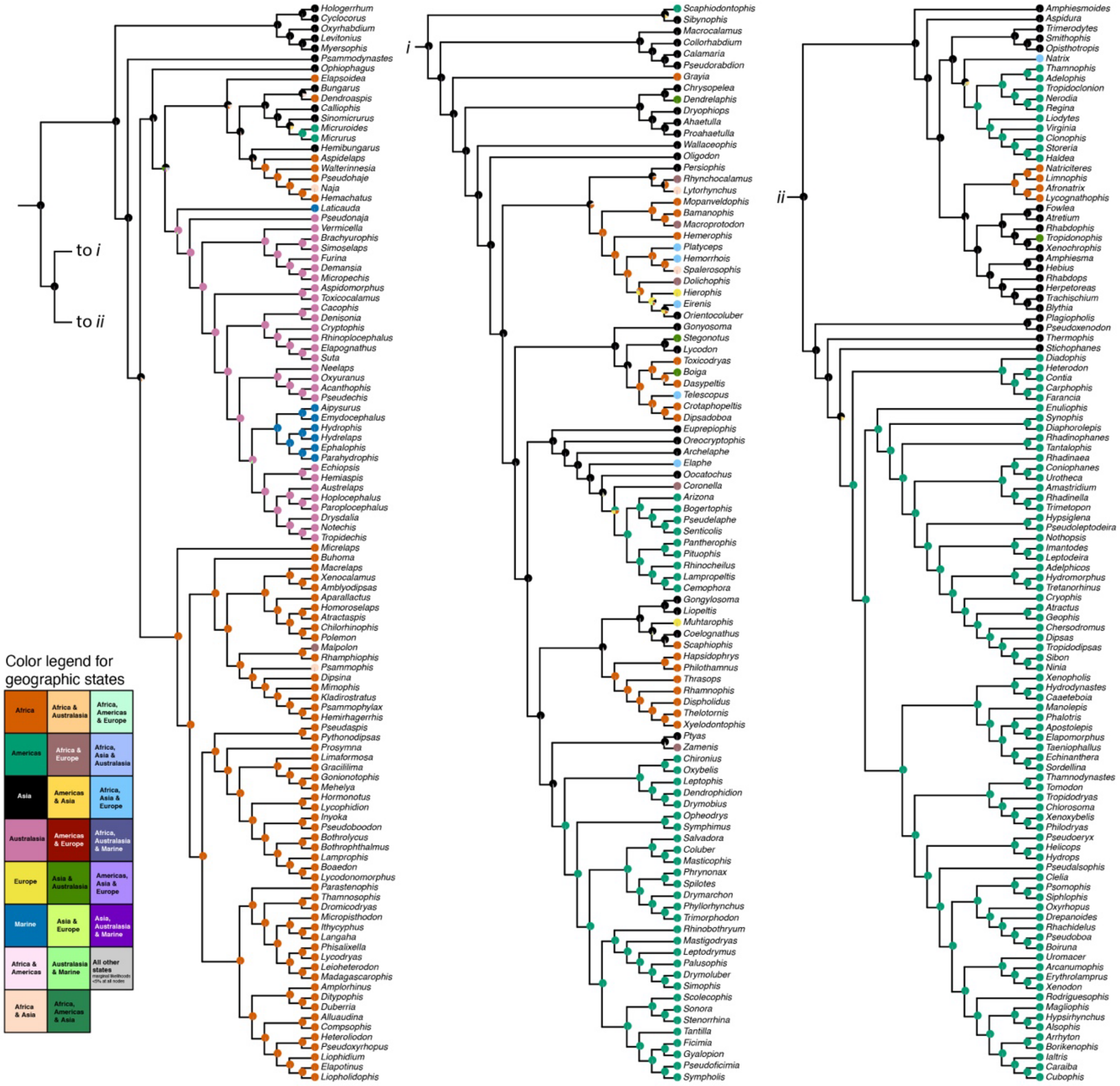


**Figure S4.** BioGeoBEARS geographic range evolution results of Elapoidea and Colubroidea under the biogeographic model "DIVALIKE+J". Node pie charts depict marginal probability estimates of ancestral geographic states. Tip circles depict geographic states of extant lineages provided as input during analysis. Phylogeny used during analysis is visualized here as a cladogram.

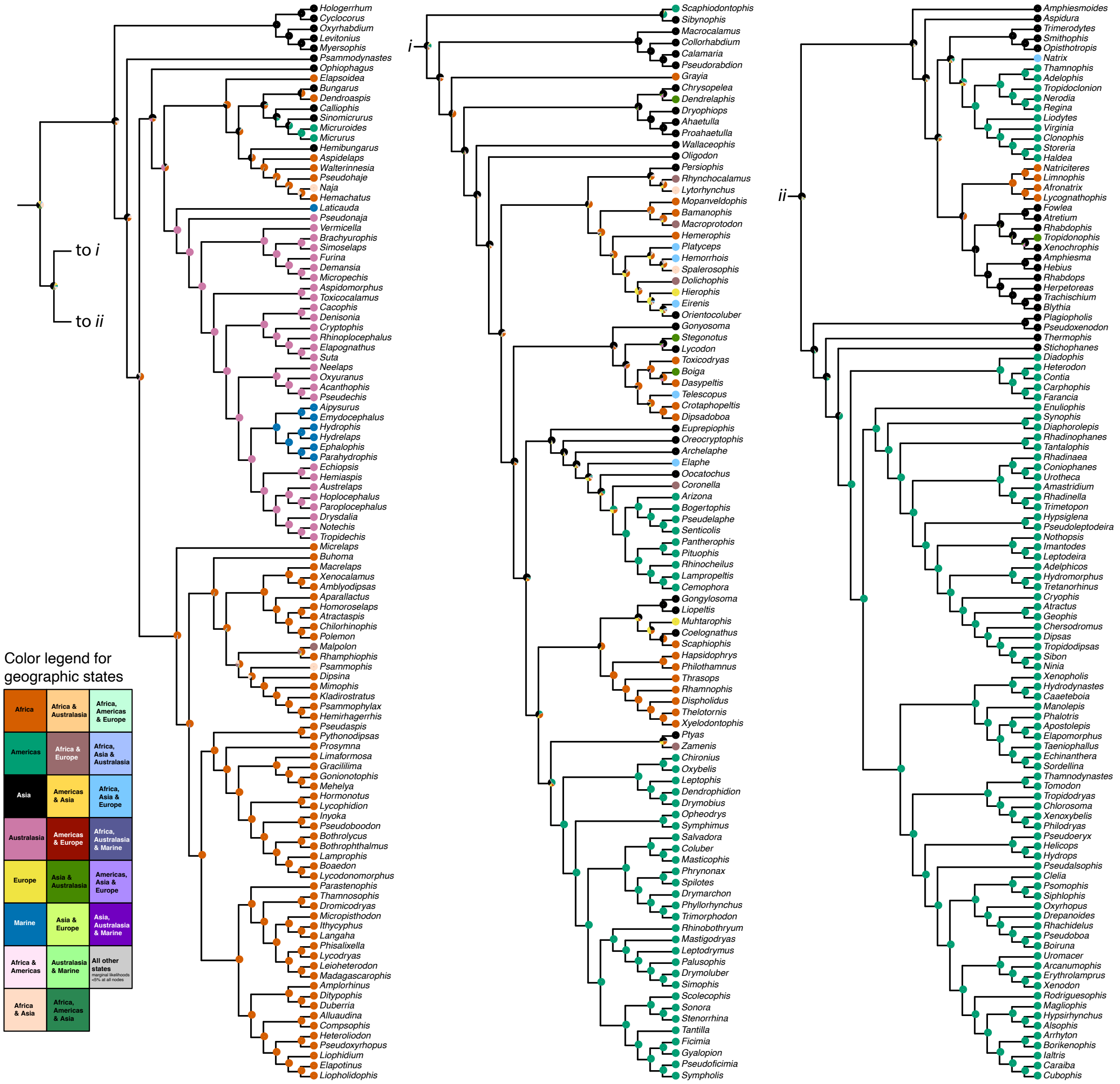


**Figure S5.** BioGeoBEARS geographic range evolution results of Elapoidea and Colubroidea under the biogeographic model "DEC+J". Node pie charts depict marginal probability estimates of ancestral geographic states. Tip circles depict geographic states of extant lineages provided as input during analysis. Phylogeny used during analysis is visualized here as a cladogram.

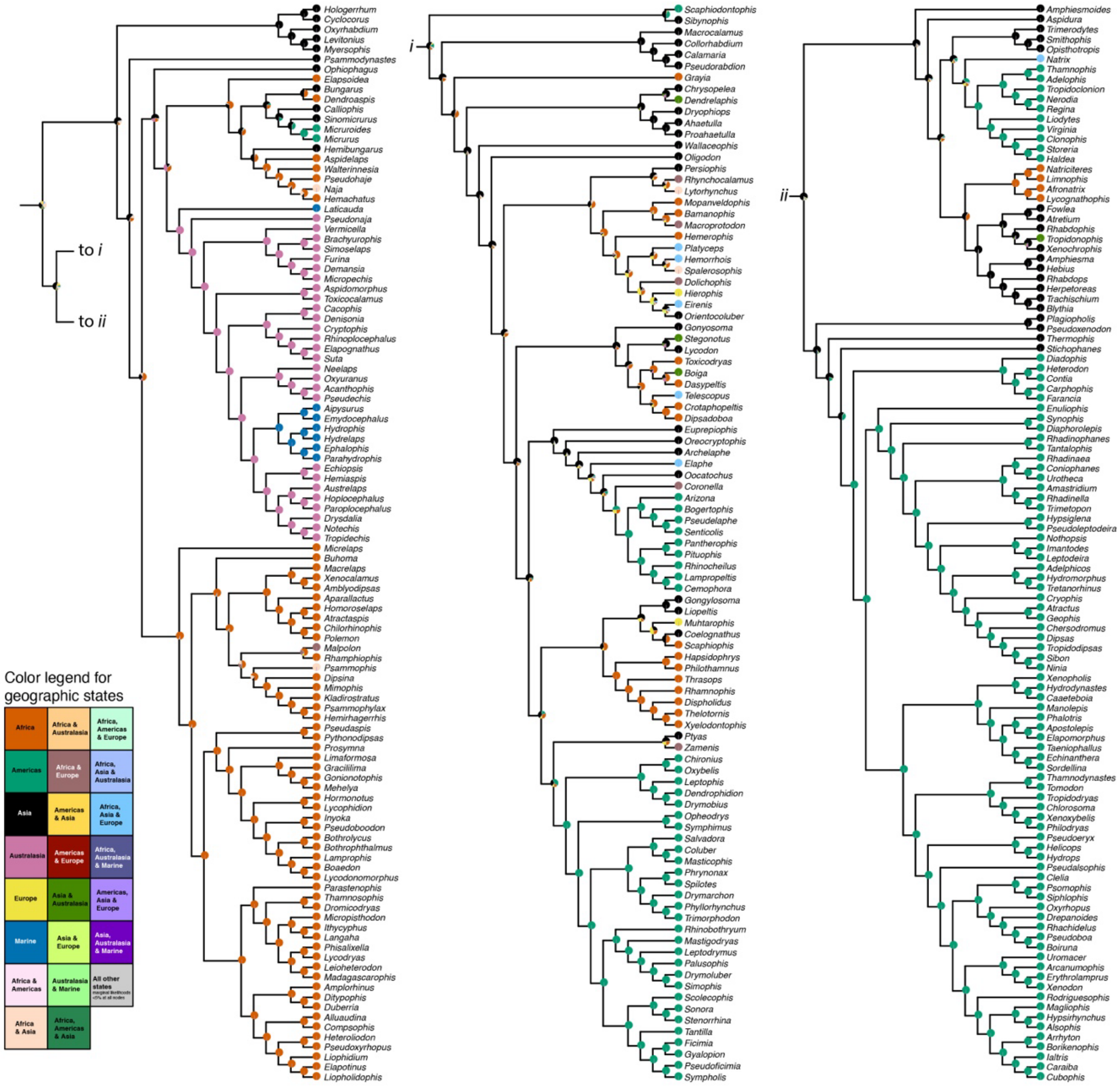


**Figure S6.** BioGeoBEARS geographic range evolution results of Elapoidea and Colubroidea under the biogeographic model “DEC”. Node pie charts depict marginal probability estimates of ancestral geographic states. Tip circles depict geographic states of extant lineages provided as input during analysis. Phylogeny used during analysis is visualized here as a cladogram.

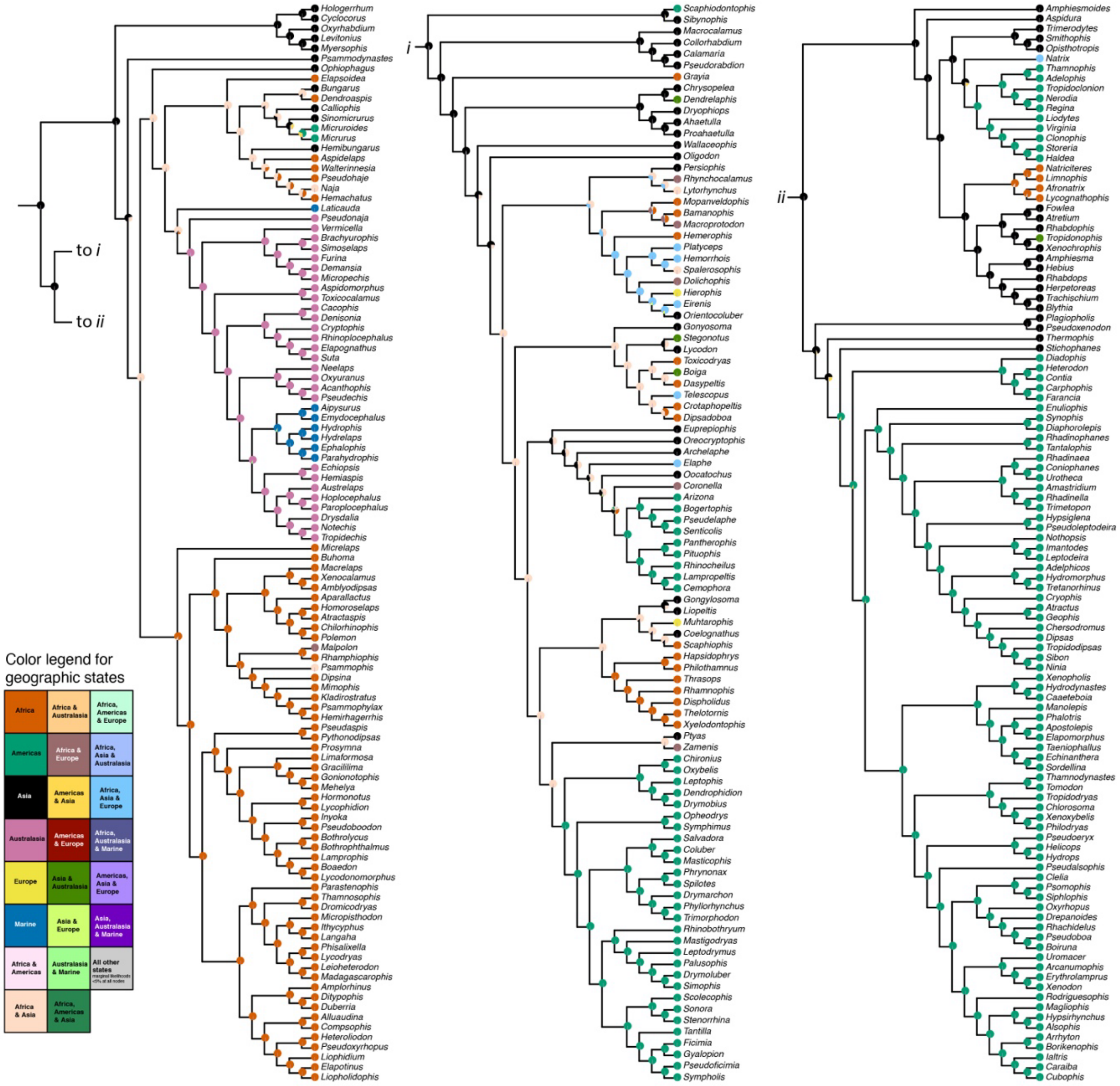


**Figure S7.** BioGeoBEARS geographic range evolution results of Elapoidea and Colubroidea under the biogeographic model "BAYAREALIKE". Node pie charts depict marginal probability estimates of ancestral geographic states. Tip circles depict geographic states of extant lineages provided as input during analysis. Phylogeny used during analysis is visualized here as a cladogram.

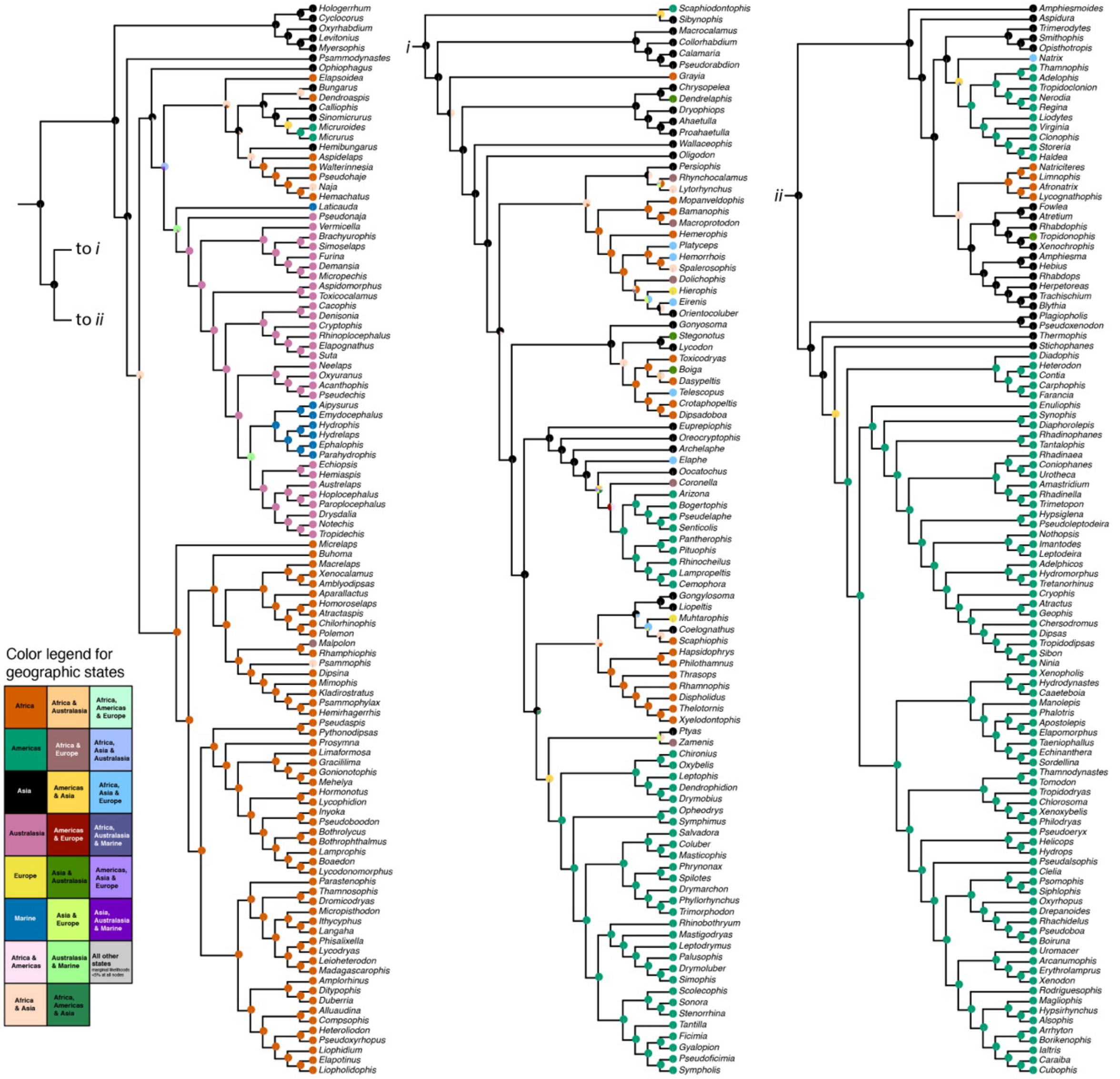


**Figure S8.** BioGeoBEARS geographic range evolution results of Elapoidea and Colubroidea under the biogeographic model "DIVALIKE". Node pie charts depict marginal probability estimates of ancestral geographic states. Tip circles depict geographic states of extant lineages provided as input during analysis. Phylogeny used during analysis is visualized here as a cladogram.

7. de Lang R. 2011 *The snakes of the Lesser Sunda Islands (Nusa Tenggara), Indonesia: A field guide to the terrestrial and semi-aquatic snakes of the Lesser Sunda Islands with identification key*. Frankfurt am Main: Edition Chimaira.

8. de Lang R. 2013 *The snakes of the Moluccas (Maluku), Indonesia: A field guide to the land and non-marine aquatic snakes of the Moluccas with identification key*. Frankfurt am Main: Edition Chimaira.

9. de Lang R, Vogel G. 2005 *The snakes of Sulawesi: A field guide to the land snakes of Sulawesi with identification keys*. Frankfurt am Main: Edition Chimaira.

10. Kurnaz M. 2020 Species list of amphibians and reptiles from Turkey. *J. Anim. Divers.* **2**, 10–32. (doi:10.29252/JAD.2020.2.4.2)

11. Latifi M. 1991 *The snakes of Iran*. Oxford, Ohio: Society for the Study of Amphibians and Reptiles (SSAR).

12. Untergasser A, Cutcutache I, Koressaar T, Ye J, Faircloth BC, Remm M, Rozen SG. 2012 Primer3—new capabilities and interfaces. *Nucleic Acids Res.* **40**, e115–e115. (doi:10.1093/nar/gks596)

13. Burbrink FT, Lawson R, Slowinski JB. 2000 Mitochondrial DNA phylogeography of the polytypic North American Rat Snake (*Elaphe obsoleta*): A critique of the subspecies concept. *Evolution* **54**, 2107–2118. (doi:10.1554/0014-3820(2000)054[2107:MDPOTP]2.0.CO;2)

14. Weinell JL, Paluh DJ, Siler CD, Brown RM. 2020 A new, miniaturized genus and species of snake (Cyclocoridae) from the Philippines. *Copeia* **108**, 144–150. (doi:10.1643/CH2020110)

15. Lawson R, Slowinski JB, Crother BI, Burbrink FT. 2005 Phylogeny of the Colubroidea (Serpentes): New evidence from mitochondrial and nuclear genes. *Mol. Phylogenet. Evol.* **37**, 581–601. (doi:10.1016/j.ympev.2005.07.016)
